# State-specific gating of salient cues by midbrain dopaminergic input to basal amygdala

**DOI:** 10.1101/687707

**Authors:** Andrew Lutas, Hakan Kucukdereli, Osama Alturkistani, Crista Carty, Arthur U. Sugden, Kayla Fernando, Veronica Diaz, Vanessa Flores-Maldonado, Mark L. Andermann

## Abstract

Basal amygdala (BA) neurons guide associative learning via acquisition of responses to stimuli that predict salient appetitive or aversive outcomes. We examined the learning- and state-dependent dynamics of BA neurons and ventral tegmental area dopamine axons that innervate BA (VTA^DA→BA^) using two-photon imaging and photometry in behaving mice. BA neurons did not respond to arbitrary visual stimuli, but acquired responses to stimuli that predicted either rewards or punishments. Most VTA^DA→BA^ axons were activated by *both* rewards and punishments, and acquired responses to cues predicting these outcomes during learning. Responses to cues predicting food rewards in VTA^DA→BA^ axons and BA neurons in hungry mice were strongly attenuated following satiation, while responses to cues predicting unavoidable punishments persisted or increased. Therefore, VTA^DA→BA^ axons may provide a reinforcement signal of motivational salience that invigorates adaptive behaviors by promoting learned responses to appetitive or aversive cues in distinct, intermingled sets of BA excitatory neurons.

## Introduction

Attention and learning are interdependent processes that depend on motivational drives. For example, a hungry animal is motivated to learn about and attend to food-predicting cues^1^. In humans and animal models, the basolateral amygdala (BLA) is one of the earliest points in the flow of sensory information where encoding of a learned sensory cue strongly depends on the current value of associated outcomes, which in turn depends on motivational state^1–3^ (see also Fig. 1a). Cue-outcome associative learning involves largely separate populations of BLA excitatory neurons that are selectively activated by either appetitive or aversive outcomes^4^ and implicated in guiding approach or avoidance behaviors. How these populations acquire selective responses to specific, motivationally salient cues remains unclear. Recent studies suggest that a simple Hebbian plasticity rule alone cannot explain the acquisition of predictive cue responses in BLA neurons *in vivo,* and argue that an additional reinforcement signal to BLA is necessary^5, 6^.

**Fig. 1.**
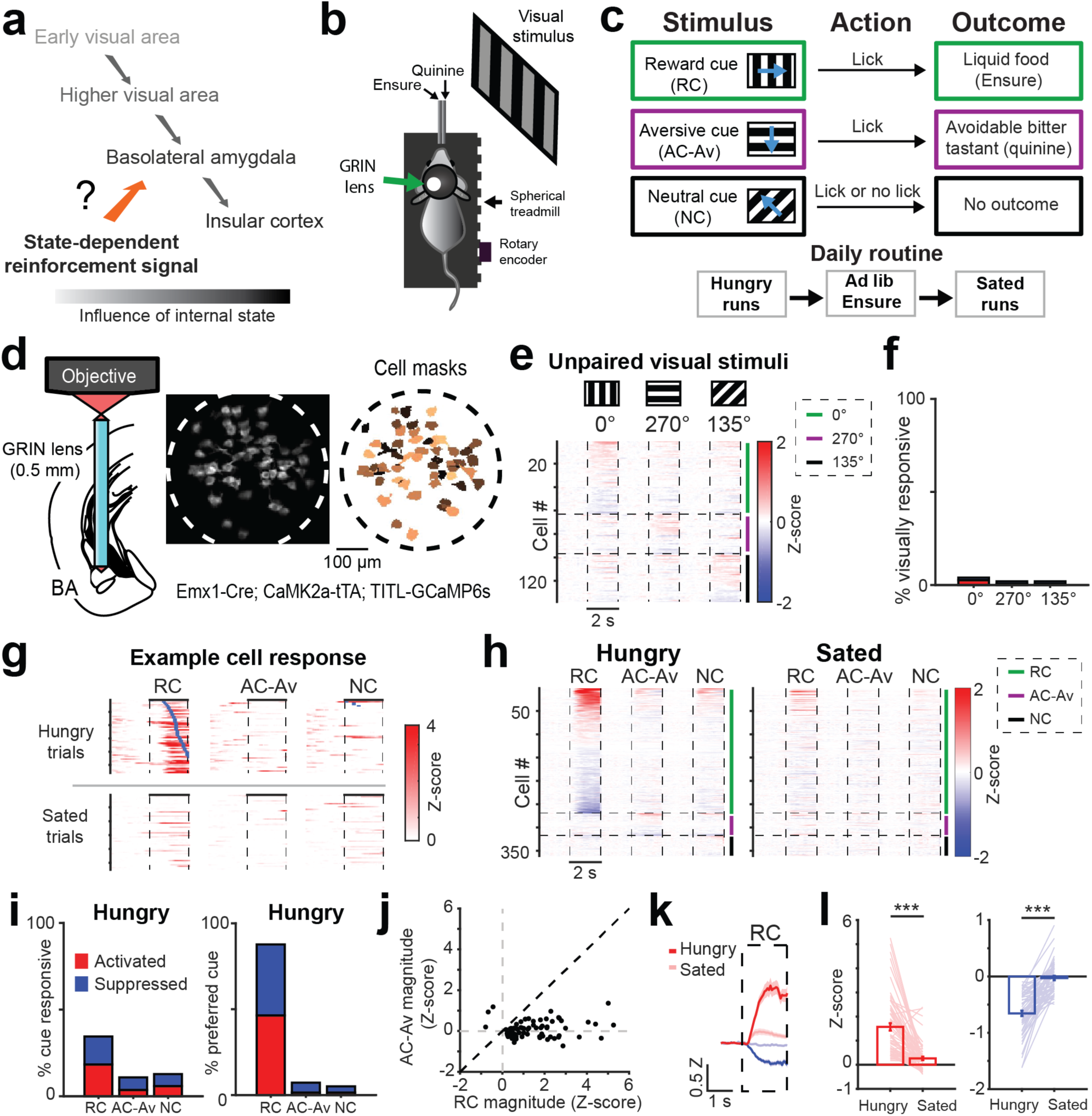
Mouse basal amygdala neurons acquire hunger-dependent responses to food-predicting cues. **a,** Visual responses along the visual pathway increasingly depend on learned motivational relevance in humans and mice. Inputs to basolateral amygdala (BLA) that relay state-specific reinforcement signals may regulate acquisition and expression of learned responses to motivationally salient cues. **b,** Schematic of head-fixed Go/NoGo visual discrimination task and imaging setup. **c,** Visual discrimination task. Mice learn that behavioral responses (licks) in the 2-s window following presentation of the 2-s reward cue (RC; oriented drifting grating) lead to liquid food delivery (Ensure). Licking following the aversive cue (AC-Av) leads to quinine delivery. This aversive outcome can be passively avoided by withholding licking. Licking following the neutral cue (NC) does not result in any outcome, regardless of action. **d,** *Left:* schematic of two-photon imaging of basal amygdala (BA) neurons using a GRIN lens (0.5 mm diameter) in transgenic mice expressing GCaMP6s in excitatory neurons. *Middle:* example field of view (ICA-based weighted cell masks, see Methods). *Right:* binarized cell masks for all active neurons, pseudocolored for visualization purposes. **e,** Heatmap with rows depicting mean responses of BA neurons (n = 137 neurons, 6 fields of view from 4 mice) to visual stimuli *prior* to associative learning, sorted by magnitude of cue response and grouped by preferred cue type for visualization. Vertical dashed lines demarcate visual stimulus onsets and offsets. Horizontal lines demarcate sorting of neurons by preferred cue (cue with the largest absolute value response). Grouping by preferred cue is also indicated by colored vertical bars to the right of the heatmap (green: 0°; purple: 270°; black: 135°). **f,** Percentage of all neurons shown in panel **e** that had a significant response to visual stimuli (see Methods; 0°: 6/137 neurons; 270°: 3/137; 135°: 3/137). **g,** Single-trial responses of an example RC-preferring neuron following associative learning. Following satiation, this neuron becomes unresponsive. Rows: trials sorted by onset of first lick (blue ticks) after visual stimulus onset. **h,** Heatmap depicting mean responses of all BA neurons (n = 360 neurons, 15 fields of view from 7 mice) during presentation of visual stimuli *after* associative learning, grouped by preferred cue type. Vertical dashed lines demarcate visual stimulus onsets and offsets. Horizontal lines demarcate sorting of neurons by preferred cue (cue with the largest absolute value response). Grouping by preferred cue is also indicated by colored vertical bars to the right of the heatmap (green: RC; purple: AC-Av; black: NC). **i,** *Left*: percentage of all neurons with significant cue responses (RC: 124/360 neurons; AC-Av: 39/360; NC: 46/360). *Right*: percentage of cue responsive BA neurons preferring (i.e., maximally responsive to) a given cue. **j,** RC vs. AC-Av cue response magnitude of activated neurons (n = 70). **k,** Average timecourse of RC responses across activated BA neurons (hungry: dark red; sated: light red; n = 66) and suppressed BA neurons (hungry: dark blue; sated: light blue; n = 58). Z: Z-score. **l,** Bars: mean RC response for activated neurons (*left,* n = 66) and suppressed neurons (*right,* n = 58). Lines: individual cell responses across hunger and satiety. *** p < 0.001, Wilcoxon sign-rank.

Dopamine is an attractive candidate teaching signal that could guide reward and aversive conditioning^7–9^ by shaping plasticity in BLA subregions including the basal amygdala (BA)^9^ and lateral amygdala (LA)^10, 11^. While pharmacological manipulations of dopamine in BLA suggest that intact dopaminergic signaling is important for associative learning^12^, the source of dopamine is unclear. Dopaminergic inputs from the ventral tegmental area (VTA) appear to selectively innervate the BA, but not the LA (see below). Lesion studies provide indirect evidence that VTA dopamine inputs to the BA (VTA^DA→BA^) are involved in learning not only of appetitive^9^ but also of aversive^8^ cue-outcome associations. While subsets of dopamine neurons within and outside of VTA are known to be activated by aversive stimuli^13–16^, it remains controversial whether VTA^DA→BA^ axons are activated by cues that predict aversive and/or rewarding outcomes^15^.

Here, we used fiber photometry and two-photon calcium imaging to examine cue responses of VTA^DA→BA^ axons in behaving mice and directly compare these responses with those of excitatory target neurons in BA. We show that, in contrast to VTA dopamine axons that innervate the nucleus accumbens (VTA^DA→NAc^), VTA^DA→BA^ axons are activated by motivationally salient appetitive and aversive outcomes, and become responsive to initially arbitrary visual cues following pairing with these outcomes. Such valence-independent responses were evident in individual VTA^DA→BA^ axons, suggesting that actions of VTA^DA→BA^ input onto any given BA target neuron likely occur during the presentation of both appetitive and aversive cues. Satiation attenuated VTA^DA→BA^ responses to reward cues while potentiating responses to cues predicting unavoidable punishments, suggesting a transition to a defensive state.

In contrast to VTA^DA→BA^ axonal inputs, we found that intermingled neurons throughout the anterior-posterior axis of BA selectively encode either appetitive *or* aversive cues during our task. Most neurons throughout BA expressed D1 dopamine receptors, and VTA^DA→BA^ axons were confirmed to release dopamine *in vivo*. Thus, activation of the same VTA^DA→BA^ axons by both appetitive and aversive cues and/or outcomes may open a state-dependent window for plasticity across most or all BA neurons. Via glutamate co-release onto inhibitory BA neurons, such activation may also drive mutual inhibition between intermingled subpopulations of appetitive- and aversive-coding BA neurons, restricting the set of neurons that acquire strong cue responses^17–19^ during associative learning.

## Results

### BA neurons are strongly biased to food-predicting visual stimuli in hungry mice

To assess the role of BA neurons in associative learning of cues predicting motivationally salient outcomes (e.g. food cues during hunger, threat cues during defensive states), we recorded responses of BA excitatory neurons to visual stimuli that either did or did not predict food reward delivery in hungry mice performing a Go/NoGo operant visual discrimination task^20^. This allowed direct comparison with neuronal responses upstream in visual cortex and LA^20, 21^ and downstream in insular cortex^22^ during identical task conditions (Fig. 1a).

Head-fixed, food-restricted mice were trained to lick following one visual stimulus (reward cue; RC) to obtain liquid food (Ensure), and to withhold licking following a different visual stimulus (avoidable aversive cue; AC-Av) to avoid a bitter tastant (quinine) (Fig. 1b,c). Licking following a third visual stimulus (neutral cue, NC) had no outcome. Mice learned to accurately perform this task within 1-2 weeks (Supplementary Fig. 1a). After performing the task in a hungry state during each session, mice received free access to Ensure until they voluntarily stopped licking (i.e. satiation; Fig. 1c). We then presented the same visual stimuli again and assessed changes in cue responses.

To image the activity of the same BA neurons in awake behaving mice with high sensitivity across hours and days, we performed two-photon calcium imaging through a gradient refractive index (GRIN) lens implanted above the BA of transgenic mice stably expressing GCaMP6s in BA excitatory neurons (Fig. 1d; Emx1-Cre; CaMK2a-tTA; TITL-GCaMP6s). BA neurons were not significantly responsive to visual stimuli in untrained mice (Fig. 1e,f), suggesting that BA neurons are selectively responsive to motivationally salient learned cues^3^. Indeed, following training, we observed strong responses to the RC (Fig. 1g,h). In contrast, we observed weak responses to the AC-Av, the NC, and to any visual cue following satiation (Fig. 1g,h). In hungry mice, a large fraction of BA excitatory neurons showed significant and selective responses to the RC (34%, 124/360 neurons from 7 mice across 15 imaging fields of view within BA; Fig. 1h-j, Supplementary Fig. 1b). Following satiation, cue responses were substantially attenuated (Fig. 1k,l). While arousal (pupil area and locomotion) was also reduced (Supplementary Fig. 2a,b), this did not account for the attenuation in RC responses, which persisted when considering trials matched for pupil dilation or locomotion across states^22^ (Supplementary Fig. 2c-f). Thus, learned RC responses in BA strongly depended on motivational state. We next examined possible reinforcement signals that could facilitate this state-dependent learning process (Fig. 1a).

### VTA^DA^**^→^**^BA^ axons show hunger-dependent responses to cues predicting food reward

Dopaminergic axonal inputs from VTA provide an attractive candidate reinforcement signal for promoting acquisition of responses to salient cues in BA neurons. As compared to LA, BA expresses higher levels of D1 receptors^23, 24^ (Supplementary Fig. 3a,b) and receives denser input from VTA dopamine axons (Fig. 2a; Supplementary Fig. 3c). To monitor the activity of VTA^DA→BA^ axons, we selectively expressed GCaMP6s in VTA dopamine neurons (in DAT-IRES-Cre mice) and placed an optic fiber above the BA, allowing bulk photometry recording of activity from VTA^DA→BA^ axons (Fig. 2b; Supplementary Fig. 4).

**Fig. 2.**
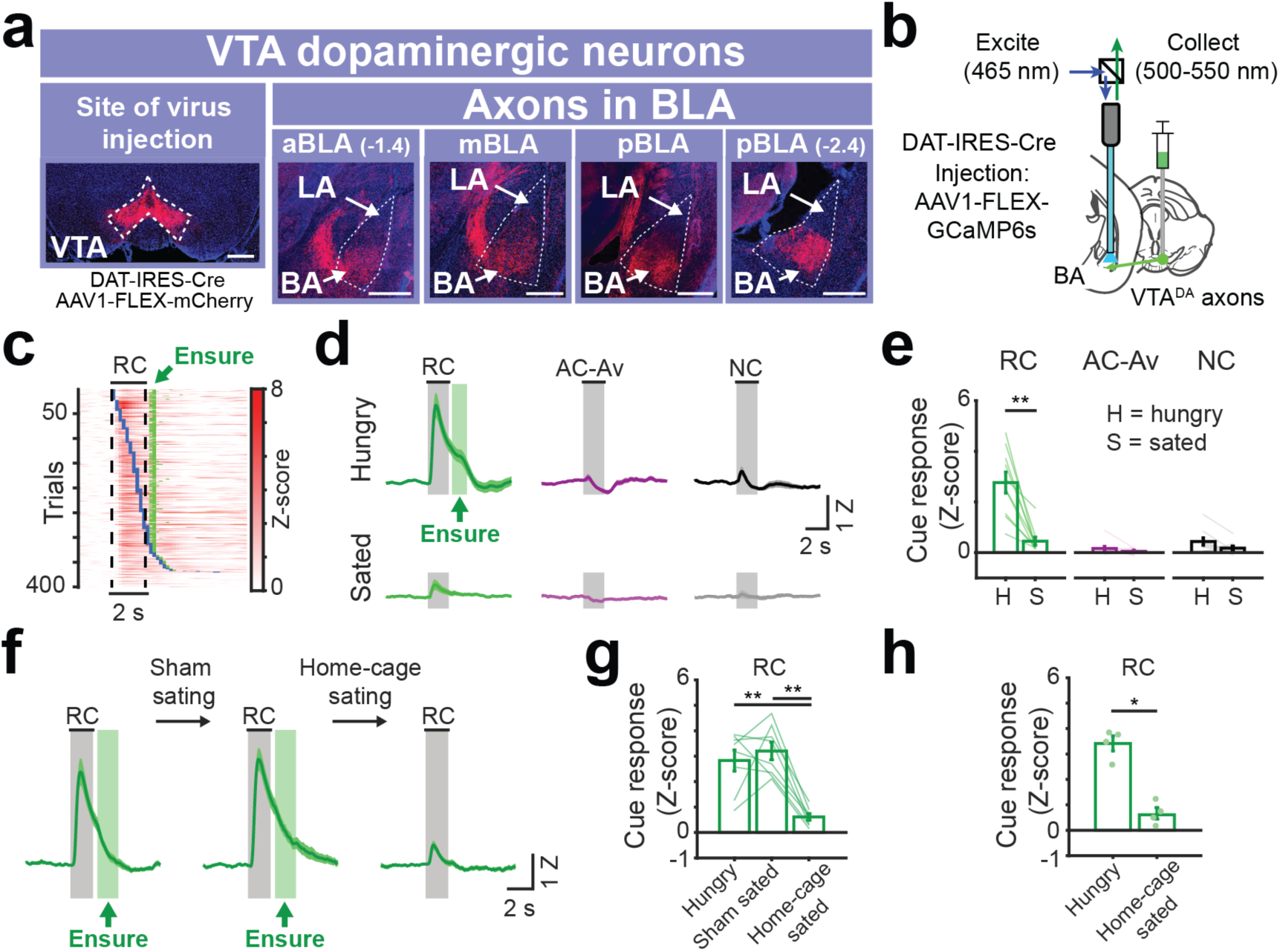
VTA dopamine axons in BA are activated by cues predicting food reward availability in hungry mice. **a,** Anterograde injection of AAV1-FLEX-mCherry in VTA of DAT-IRES-Cre mice. Within the BLA, VTA dopamine axons are strongly enriched in the basal amygdala (BA) vs. the lateral amygdala (LA). This was true in anterior, middle, as well as posterior BLA (aBLA, mBLA, and pBLA, respectively). Scale bar = 0.5 mm. **b,** Schematic of fiber photometry recordings from VTA dopamine axons in BA. **c,** Rows: all single-trial responses from 10 trained mice, sorted by onset of first lick following RC onset (blue ticks). Green ticks: time of Ensure delivery. Responses were locked to cue onset, not to motor response onset. **d,** Mean VTA^DA→BA^ cue responses in trained mice across hungry (*top*) and sated (*bottom*) states. The range of Ensure delivery times (min and max) is indicated by the green rectangle following cue offset. n = 10 mice. Error bars: s.e.m. across mice. Z: Z-score. **e,** Comparison of cue response magnitudes across states (n = 10 mice, mean ± s.e.m. across mice, ** p < 0.01, Wilcoxon sign-rank). H: hungry; S: sated. **f,** Mean VTA^DA→BA^ RC responses in hungry, sham sated, and home-cage sated mice, respectively. n = 8 mice. Error bars: s.e.m. across mice. **g,** Comparison of RC response magnitudes prior to and following sham satiation and following home-cage satiation (n = 8 mice, mean ± s.e.m. across mice, ** p < 0.01, Wilcoxon sign-rank). **h,** Comparison of RC response magnitudes following sham satiation in home cage (n = 4 mice) vs. satiation in home cage (n = 4 separate mice; mean ± s.e.m. across mice, * p < 0.05, Wilcoxon rank-sum).

Consistent with prediction error signals reported in VTA dopamine neurons^25, 26^, VTA^DA→BA^ axon activity increased to reward delivery early in training (Supplementary Fig. 5a). Once mice achieved high task performance, VTA^DA→BA^ axon responses shifted to tracking the RC (Supplementary Fig. 5b-e). RC responses were tightly locked to cue onset, and not to onset of cue-induced licking (Fig. 2c). Following satiation, RC responses were strongly attenuated (Fig. 2d-e). Sham satiation and home-cage caloric repletion experiments confirmed that this RC response attenuation was not due to incidental factors such as fatigue or stress, which might differ between early and late epochs of a given recording session (Fig. 2f-h). In contrast, the NC and AC-Av rarely elicited operant behavior in trained mice and did not evoke substantial responses in any state (Fig. 2d-e).

These findings suggested that VTA^DA→BA^ axons might signal salient outcomes and, subsequently, cues that predict salient outcomes. Supporting this hypothesis, we observed that during early training sessions with high false alarm rates, VTA^DA→BA^ axons were also activated by the quinine cue and quinine delivery (Supplementary Fig. 5f-i). While dopamine is known to be involved in amygdala plasticity during aversive associative conditioning^7, 27^, direct evidence of phasic increases in dopaminergic signaling in BA in response to reward and aversive cues and outcomes is not well established, nor is it known whether the source of dopamine during reward and aversive conditioning originates from VTA. These findings led us to consider whether VTA^DA→BA^ axons more generally display unsigned phasic responses to salient appetitive and aversive cues and outcomes rather than strictly encoding reward prediction error. However, we were unable to investigate this possibility using the AC-Av, as mice learned not to react to this cue, thereby passively avoiding quinine delivery (Supplementary Fig. 5a,b). This likely resulted in lower effective salience of the AC-Av following learning. Therefore, we next considered VTA^DA→BA^ responses to *unavoidable* aversive outcomes and associated cues.

### Cues predicting unavoidable aversive outcomes also activate VTA^DA^**^→^**^BA^ axons

If VTA^DA→BA^ axons carry information about motivational salience rather than strictly about rewards, they should also acquire responses to cues predicting highly salient aversive outcomes (Fig. 3a). To test this, mice previously trained on the task involving an aversive cue predicting avoidable quinine were subsequently trained on a modified task in which the aversive cue now predicted an *unavoidable* aversive outcome (air puff delivered to the face; Fig. 3b; task performance remained high with few false alarms, Supplementary Fig. 6a,b). Strikingly, we observed a large increase in VTA^DA→BA^ axon activity in response to air puffs, and emergence of significant responses to the aversive cue predicting this unavoidable outcome (AC-Un) within the first session (Fig. 3c, Supplementary Fig. 6c-e). Air puff responses decreased across sessions, accompanied by a concomitant increase in AC-Un responses (Supplementary Fig. 6d-f). These effects were not due to overall changes in network excitability, as RC response magnitudes remained stable throughout these sessions (Fig. 3c; Supplementary Fig. 6g). In contrast to the lack of any obvious behavioral responses to the avoidable quinine-predicting cue following training, mice developed an active avoidance behavior – blinking – to the AC-Un (Supplementary Fig. 7a-b). Taken together, these findings suggest that cue-evoked increases in activity of VTA^DA→BA^ axons reflect an *unsigned* (i.e. valence-independent) signal of the motivational salience of appetitive and aversive predicted outcomes.

**Fig. 5.**
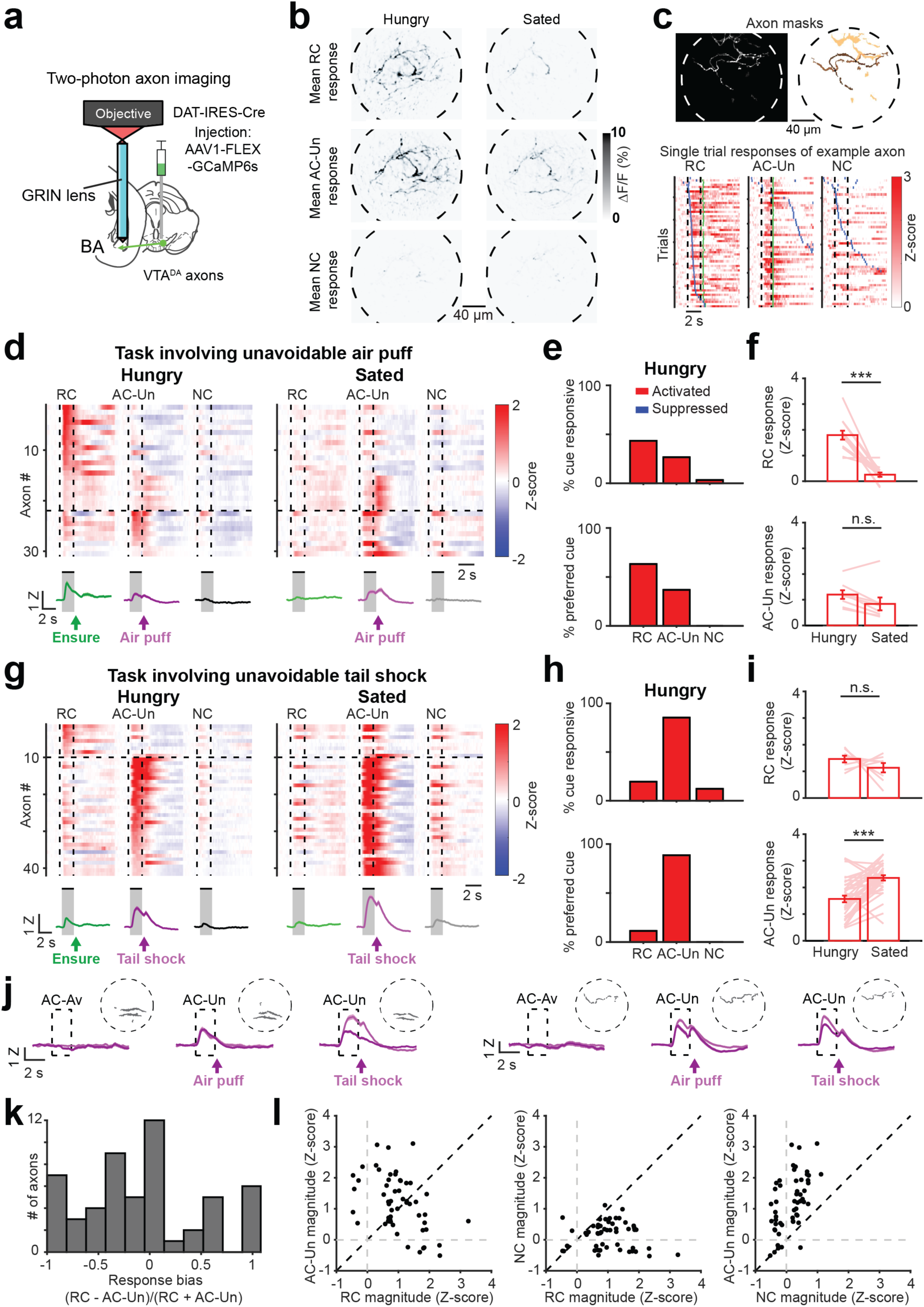
Individual VTA^DA→BA^ axons are activated by both cues predicting reward and cues predicting unavoidable aversive outcomes. **a,** Schematic depicting two-photon calcium imaging of VTA^DA→BA^ axons using a GRIN lens. **b,** Mean cue-evoked responses (fractional change in GCaMP6s fluorescence, averaged across 2-s stimulus presentation) across a single session for an example field of view during hungry trials (*left column*) or sated trials (*right column*), from the task involving cues predicting unavoidable aversive air puff (AC-Un). Note that individual axons are activated by both RC and AC-Un, and that AC-Un responses in most axons persist during sated trials. See also Supplementary Movie 1. **c,** *Top left:* maximum intensity projection across ICA-based weighted axon masks from an example field of view. *Top right:* masks for 3 responsive axons, thresholded and pseudocolored for visualization purposes. *Bottom:* Single-trial responses of an example VTA^DA→BA^ axon following associative learning. This axon is responsive to both the reward cue (RC) and the aversive cue predicting unavoidable air puff (AC-Un). Rows: trials sorted by onset of first lick after visual stimulus onset (blue ticks). Green ticks: Ensure delivery during RC trials or air puff delivery during AC-Un trials. **d,** *Top*: mean response of individual axons (rows) to presentation of the RC, AC-Un, and neutral cue (NC) (n = 30 axons, 6 fields of view from 4 mice). *Bottom:* mean cue response timecourse across all axons. Error bars: s.e.m. across axons. Z: Z-score. Data are from task in which AC-Un predicts air puff. Vertical dashed lines demarcate visual stimulus onsets and offsets. Horizontal lines demarcate sorting of axons by preferred cue (cue with the largest absolute value response). **e,** *Top*: percent of all axons with significant cue responses (RC: 13/30 axons; AC-Un: 8/30; NC: 1/30). *Bottom*: percent of cue responsive axons preferring a given cue. All data from recordings during the hungry state. **f,** *Top*: mean RC response magnitude for significantly activated axons. Lines: individual axon responses across hunger and satiety. *Bottom*: mean AC-Un response for activated axons. *** p < 0.001, n.s., not significant, Wilcoxon sign-rank. **g,** *Top*: mean cue responses of individual axons (rows) (n = 41 axons, 7 fields of view from 4 mice). *Bottom:* mean cue response timecourse across all axons. Error bars: s.e.m. across axons. Z: Z-score. Data are from task in which AC-Un predicts tail shock. **h,** *Top*: percent of all axons with significant cue responses (RC: 8/41 axons; AC-Un: 35/41; NC: 5/41). *Bottom*: percent of cue responsive axons preferring a given cue. **i,** *Top*: mean RC response for significantly activated axons. Lines: individual axon responses across hunger and satiety. *Bottom*: mean AC-Un response for activated axons. *** p < 0.001, n.s., not significant, Wilcoxon sign-rank. **j,** Mean response timecourses across aversive cue trials for two axons (recorded in separate mice) that were tracked across sessions spanning 18 days. Single-session timecourses are shown following training on successive versions of the task in which the same aversive cue predicted avoidable quinine, then unavoidable air puff, and then unavoidable tail shock. Traces from hungry runs (dark purple) and sated runs (light purple) are shown. Note that both axons were not responsive to AC-Av but then acquired responses to both the AC-Un paired with air puff and to the AC-Un paired with tail shock. Error bars: s.e.m. across trials (on average, 50 trials/session). Dashed circles: perimeter of field of view with ICA-based single-axon mask (thresholded). **k,** Histogram of cue response bias of VTA^DA→BA^ axons significantly activated by either the RC and/or the AC-Un. Note the number of axons with similar magnitude responses to both cues (i.e. values near zero). **l,** Scatter plot of magnitudes of RC vs. AC-Un responses (*left;* Pearson’s r: −0.46, p < 0.001), RC vs. NC responses (*middle;* not significantly correlated), and NC *vs.* AC-Un responses (*right;* Pearson’s r: 0.56, p < 0.001). Data in **k** and **l** were combined across tasks involving air puff (**d**) and tail shock (**g**).

**Fig. 7.**
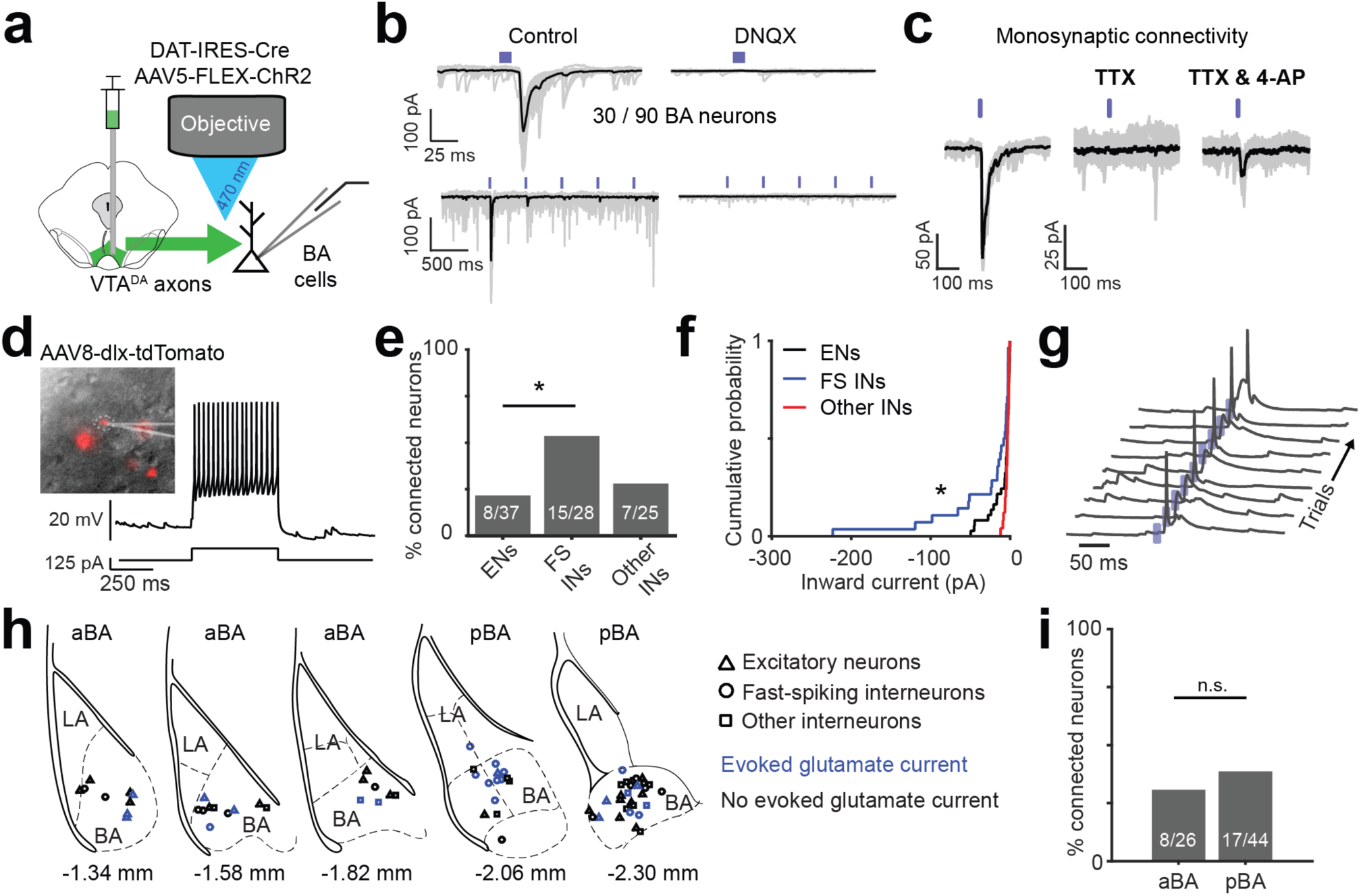
VTA^DA→BA^ axons release glutamate. **a,** Schematic of channelrhodopsin (ChR2)-assisted circuit mapping of connectivity between VTA dopaminergic axons and BA neurons. **b,** *Top left:* example recording of ChR2-evoked inward synaptic current (−70 mV holding potential). *Bottom left:* repeated stimulation at 2 Hz resulted in strong short-term depression. *Right:* evoked inward currents were blocked by a glutamate receptor antagonist (DNQX, 10 μM). **c,** Example experiment confirming monosynaptic connectivity by blocking action potentials with a voltage-gated sodium channel blocker, tetrodotoxin (TTX; 1 μM), followed by addition of a potassium channel blocker, 4-aminopyridine (4-AP; 100 μM), to restore evoked synaptic release. **d,** Identification of inhibitory interneurons using a viral strategy involving infection with AAV8-*dlx*-tdTomato (see Methods) and electrophysiological characterization. *Left*: *dlx*-tdTomato positive neuron. *Right*: current clamp recording with somatic current injection (125 pA) in a typical fast-spiking interneuron. **e,** Percentage of connected neurons, separated by neuronal class. Fast-spiking, putative inhibitory interneurons (FS INs) were more likely than excitatory neurons (ENs) or other non-fast spiking interneurons (Other INs) to evoke glutamatergic currents (n = 90 neurons from 10 mice, * p < 0.05, binomial proportion test, Bonferroni corrected). **f,** Cumulative probability distributions show that VTA^DA→BA^ axon-evoked glutamatergic currents were larger in FS INs than in ENs. * p < 0.05, Komolgorov-Smirnov test. **g,** ChR2-evoked synaptic potentials were sufficient to trigger somatic action potentials in FS INs. Traces: consecutive single trials. Blue ticks: blue light pulses. **h,** Locations of recorded neurons for which we had low magnification images of recording pipette location (70/90 neurons). Neuronal class markers: ENs = triangles, FS INs = circles, Other INs = squares. Blue: neurons with light-evoked glutamatergic currents. Sections used for comparison of anterior BA (aBA) and posterior BA (pBA) in panel **i** are indicated. Location relative to Bregma based on Paxinos and Franklin’s mouse atlas (4^th^ edition) is indicated below each section. **i,** Percentage of connected neurons, separated by aBA vs. pBA. (n = 70 neurons for which we had anatomical location from 10 mice, n.s. = not significant, binomial proportion test).

The relative magnitude of VTA^DA→BA^ axon responses to appetitive vs. aversive cues changed depending on motivational state. Satiation led to a decrease in RC responses in VTA^DA→BA^ axons and an *increase* in AC-Un responses (Fig. 3d-e), resulting in a shift in cue response preference from reward cues to aversive cues (Fig. 3f). These findings suggest a shift from a reward-seeking state to a defensive state that results in a decrease in the relative motivational salience of food-predicting vs. punishment-predicting cues. Accordingly, sated mice exhibited a persistent defensive behavior not observed in hungry mice – sustained, partial closure of the eye ipsilateral to the air puff (contralateral to the visual stimulus; Supplementary Fig. 7c-d). We next addressed whether these VTA^DA→BA^ axon responses were specific to air puff-predicting cues or whether they generalized to a different aversive outcome eliciting a distinct avoidance behavior. We replaced air puff delivery (which elicited eye closure) with a mild unavoidable tail shock that elicited increased locomotion, possibly as part of an escape response (Supplementary Fig. 6a and 7e,f; n = 6 mice previously trained on the quinine/air puff tasks and 2 newly trained mice; task performance remained high with few false alarms, Supplementary Fig. 6b). As with air puff delivery, tail shocks increased activity of VTA^DA→BA^ axons early in training (Supplementary Fig. 6c,d; Supplementary Fig. 7g), as did tail shock-predicting cues (AC-Un) after one day of training (Fig. 3g,h; Supplementary Fig. 6e). As with cues predicting air puff, satiation resulted in significantly elevated responses to tail shock-predicting cues (Fig. 3h) and a shift in response preference towards these cues (Fig. 3i). Control recordings from VTA^DA→BA^ axons expressing GFP confirmed that cue-evoked responses were not generated or influenced by motion artifacts (Supplementary Fig. 7h). Therefore, despite differences in *behavioral* responses to aversive cues that predict air puff or tail shock, both these cues evoked similar hunger-dependent responses in VTA^DA→BA^ axons.

**Fig. 6.**
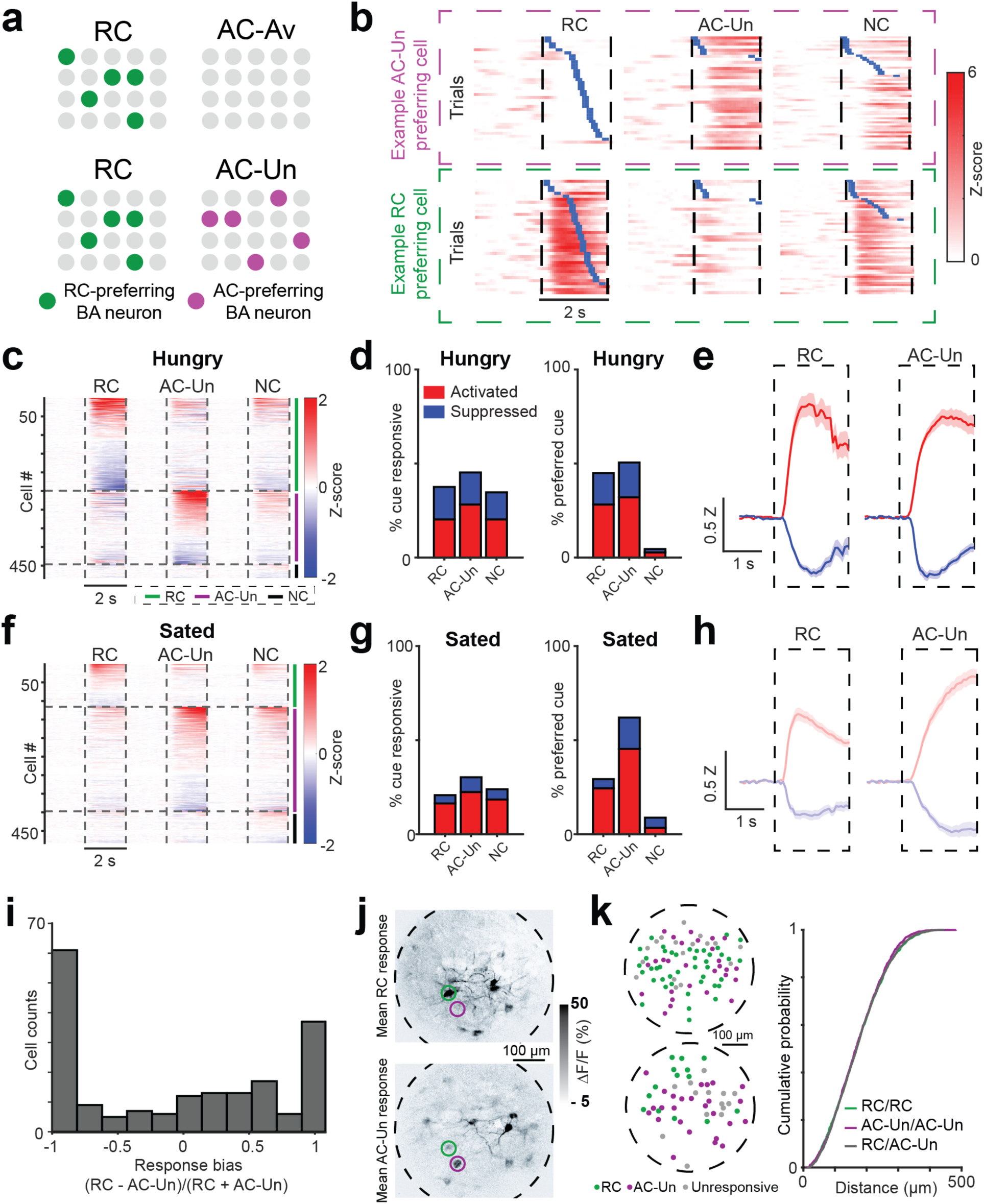
Distinct, intermingled BA neurons acquire responses to cues predicting either reward or unavoidable aversive outcomes. **a,** *Top row:* schematic depicting results from Fig. 1, in which certain basal amygdala (BA) excitatory neurons became responsive to the RC (*left*) but not to the aversive cue predicting passively avoidable quinine outcome (AC-Av, *right*). *Bottom row:* for the modified task involving cues predicting unavoidable aversive outcomes (AC-Un, *right*) as well as an RC (*left*), we asked whether certain BA neurons responded mainly to the RC (green), to the AC-Un (purple), or to both. **b,** Example BA neurons that are preferentially activated by either the AC-Un (*top)* or the RC (*bottom*). Trials are sorted by the onset of licking (blue ticks). Vertical dashed lines demarcate visual stimulus onsets and offsets. **c,** Heatmap showing mean cue response timecourses of all recorded BA neurons (rows) during hungry sessions, sorted by response magnitude and clustered by preferred cue (n = 482 neurons, 9 fields of view from 4 mice), for the task involving unavoidable tail shock. Vertical dashed lines demarcate visual stimulus onsets and offsets. Horizontal lines demarcate sorting of axons by preferred cue. **d,** *Left*: percent of BA neurons with significant cue responses during recordings in hungry mice (RC: 181/482 neurons; AC-Un: 218/482; NC: 167/482). *Right*: percent of cue responsive BA neurons preferring a given cue. **e,** Mean response timecourses across neurons that were significantly activated (red) or suppressed (blue) by the RC or the AC-Un during hungry sessions. Error bars: s.e.m. across neurons. Z: Z-score. **f,** Heatmap showing mean cue response timecourses of all recorded BA neurons (rows) during sated sessions, sorted by response magnitude and clustered by preferred cue (n = 482 neurons, 9 fields of view from 4 mice), for the task involving unavoidable tail shock. **g,** *Left*: percent of BA neurons with significant cue responses during recordings in sated mice (RC: 100/482 neurons; AC-Un: 146/482; NC: 115/482). *Right*: percent of cue responsive BA neurons preferring each cue. **h,** Mean response timecourses across neurons that were significantly activated (red) or suppressed (blue) by the RC or the AC-Un during sated sessions. Error bars: s.e.m. across neurons. Z: Z-score. **i,** Response bias of neurons activated by RC and/or AC-Un. Note the paucity of neurons with equal response magnitudes to both cues (i.e. bias value near zero) as compared to VTA^DA→BA^ axons (Fig. 5k). **j,** Mean cue-evoked responses (fractional change in GCaMP6s fluorescence) across a single session for an example field of view. *Top:* RC response. *Bottom:* AC-Un response. Even neighboring neurons could show selective responses to the AC-Un (purple circle) or to the RC (green circle). **k,** *Left:* two example fields of view showing centroids of BA neurons (green: RC responsive, purple: AC-Un responsive; gray: unresponsive). *Right*: cumulative probability distributions show that distances between neurons preferring the same cue (i.e. both preferring RC, ‘RC/RC’, or both preferring AC-Un, ‘AC-Un/AC-Un’) were not different than neurons preferring opposite cue types (RC/AC-Un) (Komolgorov-Smirnov test). Note that, given the higher resolution of two-photon imaging compared to one-photon imaging, we have high confidence in the estimated distances between cells and in the lack of cross-contamination of signals from nearby cells.

Additional evidence supported the notion that VTA^DA→BA^ axons may signal the motivational salience of cues predicting salient outcomes associated with active motor strategies. Specifically, while visual cues predicting *unavoidable* tail shock drove strong VTA^DA→BA^ axon responses following training (as well as delayed, cue-induced locomotor behavior), similar responses were not observed following training in experiments where the same visual stimuli predicted passively avoidable tail shocks (Fig. 3j-l; consistent with the lack of strong responses to cues predicting passively avoidable quinine, Fig. 2d-e).

The activation of VTA^DA→BA^ axons could represent a surprise signal in response to any violation of expectations^28^ (e.g. unexpected occurrence of a salient cue or unexpected omission of a strongly expected outcome). However, we did not observe any increase in activity following omissions of either tail shocks or rewards in well-trained mice (Supplementary Fig. 8). We sometimes observed a decrease in activity following either omitted aversive or appetitive outcomes, but only when these omission trials were preceded by a trial involving delivery of that same outcome (Supplementary Fig. 8), consistent with a negative prediction error signal. Our findings support the hypothesis that cue responses in VTA^DA→BA^ axons constitute signals of predicted motivational salience of outcomes requiring active motor behavior, rather than signals related to surprise or involved in planning of specific behavioral responses.

### Comparisons of recordings of dopaminergic activity in subregions of BA and in NAc

Recent studies have argued for functional organization of BA neurons, with enriched incidence of aversive and appetitive responses in anterior (aBA) and posterior BA (pBA), respectively^29, 30^. Thus, one might expect larger responses to the AC-Un and its associated punishment in VTA^DA→BA^ axons in aBA and larger responses to the RC and its associated reward in VTA^DA→BA^ axons in pBA^29^. By sorting based on recording location along the anterior-posterior axis (Supplementary Fig. 4a), we found that VTA^DA→BA^ axons in aBA and pBA were activated by both the RC and AC-Un (Supplementary Fig. 9a-d), similar to observations in BA cell bodies (see below and ^30^). Notably, AC-Un responses of VTA^DA→BA^ axons were not significantly different between the aBA and pBA. However, RC responses of VTA^DA→BA^ axons were significantly larger in aBA vs. pBA, contrary to expectations^29^ (but see ^31^ for evidence of selectivity for positive outcomes in aBA) (Supplementary Fig. 9a-f). We did not observe differences between aBA and pBA in the magnitude of responses to reward delivery, air puff, or tail shock (Supplementary Fig. 9e-h). These data suggest that, apart from some differences in RC response magnitude following training, VTA^DA→BA^ axons targeting aBA or pBA exhibit similar response properties.

We next sought to directly compare responses of VTA^DA→BA^ axons with those of VTA axons in the nucleus accumbens (VTA^DA→NAc^), which are known to exhibit classic reward prediction error signals^32^ and which do not strongly collateralize to the BA^33^. First, we simultaneously recorded ipsilateral calcium signals from VTA^DA→NAc^ and VTA^DA→BA^ axons (Fig. 4a,b and Supplementary Fig. 4c). We found significant RC-evoked increases in activity in both VTA^DA→NAc^ and VTA^DA→BA^ axons (Fig. 4b-d). In contrast, while the AC-Un evoked an increase in activity in VTA^DA→BA^ axons, it evoked a *decrease* in activity in VTA^DA→NAc^ axons (Fig. 4b,c), consistent with previous studies indicating that VTA dopamine neurons signal the predicted value of rewards and reward-prediction errors^26, 32^.

**Fig. 3.**
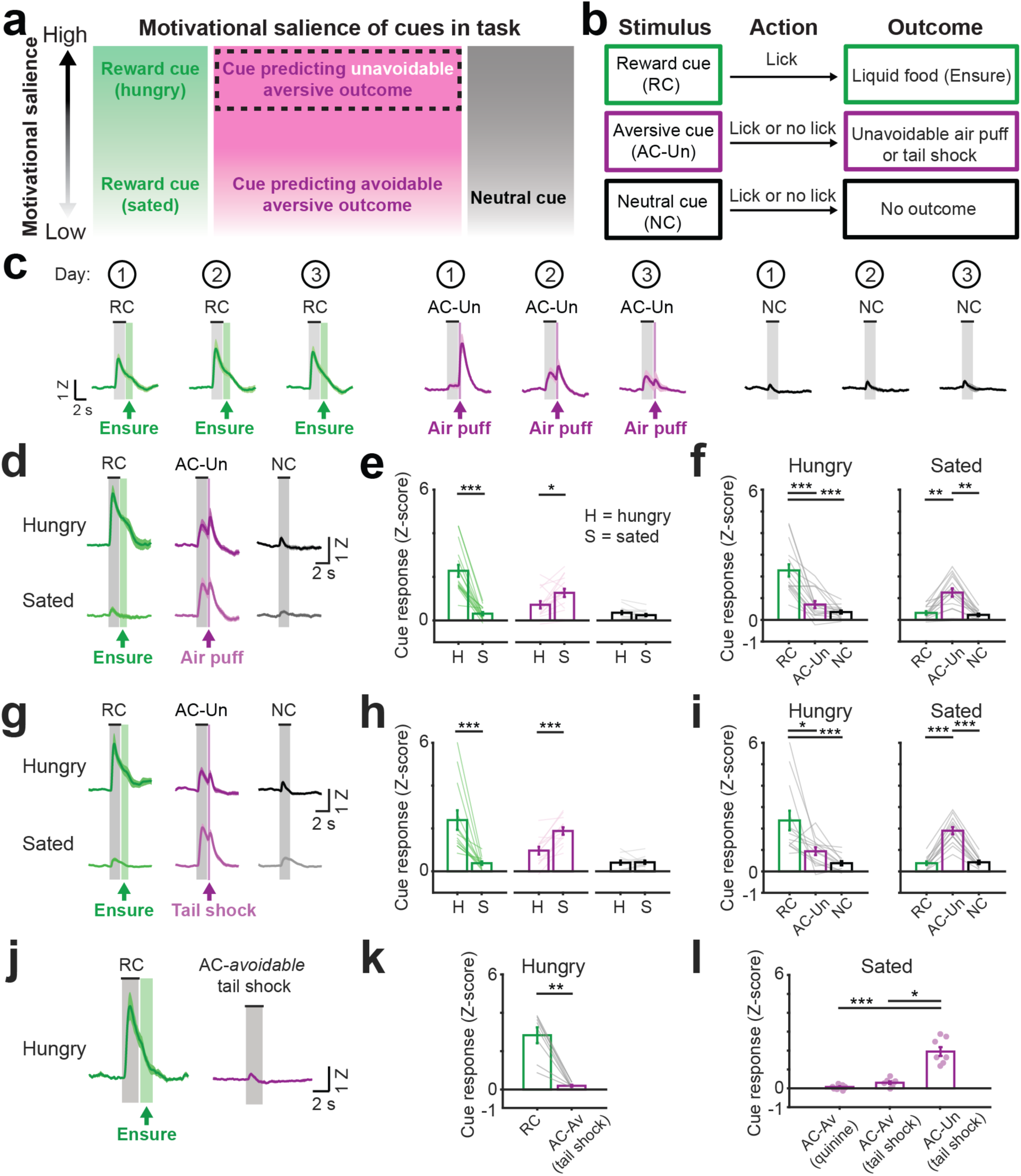
VTA dopamine axons in BA are activated by cues predicting unavoidable aversive outcomes. **a,** Diagram of relative motivational salience of each cue. We hypothesized that cues predicting unavoidable aversive outcomes would have higher motivational salience than cues predicting a passively avoidable aversive outcome or no outcome. **b,** Modified task with aversive cue (AC-Un) predicting an *unavoidable* aversive outcome (air puff or tail shock). **c,** Mean cue response timecourses of VTA^DA→BA^ axons for each of the three first sessions (numbers within circles at top) following introduction of the AC-Un predicting unavoidable air puff delivery. Mice were previously trained on the task involving *avoidable* aversive cues (AC-Av). Error bars: s.e.m. across 10 mice. Z: Z-score. **d,** Mean VTA^DA→BA^ cue responses in hungry and sated mice (combined across Days 2 and 3 from **c**) following acquisition of responses to cues predicting unavoidable aversive air puff. n = 10 mice. Error bars: s.e.m. across 16 sessions. **e,** Comparison of cue response magnitudes across states (mean ± s.e.m. across 16 sessions from 10 mice, *** p < 0.001, * p < 0.05, Wilcoxon sign-rank). H: hungry; S: sated. **f,** Comparison of cue response magnitudes within hunger state and within sated state (same data as in **e**, n = 10 mice, mean ± s.e.m. across 16 sessions, ** p < 0.01, *** p < 0.001, Kruskal-Wallis, Bonferroni corrected post-hoc comparisons). **g,** Mean VTA^DA→BA^ cue responses in hungry and sated mice (combined across Days 2 and 3 following introduction of tail shocks) after acquisition of cues predicting unavoidable tail shock. n = 8 mice. Error bars: s.e.m. across 13 sessions. **h,** Comparison of responses to the same cue across states (mean ± s.e.m. across 13 sessions from 8 mice, *** p < 0.001, Wilcoxon sign-rank). **i,** Comparison of response magnitude within hunger state and within sated state (same data as in **h**, mean ± s.e.m. across 13 sessions from 8 mice, * p < 0.05, *** p < 0.001, Kruskal-Wallis, Bonferroni corrected post-hoc comparisons). **j,** Mean VTA^DA→BA^ cue responses following training on a task involving *avoidable* tail shock in hungry mice. Error bars: s.e.m. across 8 mice. Z: Z-score. **k,** Comparison of cue response magnitudes following training on a task involving reward-predicting and avoidable tail shock-predicting cues (AC-Av tail shock). Mean ± s.e.m. across 8 mice, ** p < 0.01, Wilcoxon sign-rank. **l,** Comparison of cue response magnitudes following training on a task involving either avoidable quinine (AC-Av quinine, n = 10 mice), avoidable tail shock (AC-Av tail shock, n = 8 mice), or unavoidable tail shock (AC-Un tail shock, n = 8 mice). *** p < 0.001, * p < 0.05, Kruskal-Wallis, Bonferroni corrected post-hoc comparisons

Previous studies of dopamine dynamics in NAc and BA involve measurements of dopamine release rather than dopaminergic axon activity. To assess whether these measures yield similar results, we used a similar fiber photometry strategy to simultaneously record dopamine dynamics *in vivo* in BA and NAc using a genetically-encoded fluorescent dopamine sensor (dLight1.1^34^; Supplementary Fig. 4d). We found that dopamine levels increased in BA during the RC and AC-Un (Fig. 4e-f), with similar dynamics to VTA^DA→NAc^ axonal calcium activity. In contrast, dopamine levels in NAc *decreased* in response to the AC-Un, consistent with our calcium activity recordings and with prior dLight1.1 recordings^34^. Notably, as with axon calcium activity recordings (Fig. 4d), we observed a tight correlation in the trial-to-trial increases in dopamine levels in BA and NAc during RC trials (Fig. 4g; r = 0.57), and a weaker correlation during AC-Un trials (Fig. 4g; r = 0.12), suggesting that common inputs to VTA^DA→BA^ and VTA^DA→NAc^ neurons may drive these correlations in dopamine release.

These data suggest that our earlier findings regarding distinct coding of cues in VTA dopaminergic axons in BA vs. NAc (Figs. 2-4) do not reflect considerations specific to calcium recordings, but instead correlate with dopamine release from VTA^DA→BA^ during salient cue presentation. To confirm that dopamine is indeed released from VTA^DA→BA^ axons *in vivo*, we expressed a red-shifted excitatory opsin, Chrimson^35^, in VTA dopamine neurons and recorded dopamine levels in BA using dLight1.1 (Fig. 4h; Supplementary Fig. 4e). Stimulation of Chrimson-expressing VTA axon terminals in BA (2 mW or 5 mW, 20 Hz) drove dLight1.1 responses of roughly similar magnitude to those evoked by salient cues (Fig. 4i), supporting the notion that VTA^DA→BA^ axons provide a major source of dopamine release in BA.

### Individual VTA^DA^**^→^**^BA^ axons are activated by both cues that predict reward and cues that predict unavoidable aversive outcomes

Our bulk fiber photometry recordings showed that the average activity across many VTA^DA→BA^ axons was increased upon presentation of both the RC and the AC-Un. This response profile could be due to averaging across functionally distinct VTA^DA→BA^ axons activated by either the RC or AC-Un. Alternatively, individual VTA^DA→BA^ axons that give rise to this bulk photometry signal could respond to both the RC and AC-Un. We therefore imaged individual dopaminergic axons (3-10 axons per ∼200 μm diameter field of view) in the BA of behaving mice using two-photon calcium imaging^36^ via a GRIN lens (Fig. 5a; Supplementary Fig. 10a). In the task involving an RC, a NC, and a cue predicting *passively avoidable* quinine, we found that individual VTA^DA→BA^ axons were only responsive to the motivationally salient RC (Supplementary Fig. 10b-d). Further, the mean response time course across all VTA^DA→BA^ axons was similar to that obtained using bulk fiber photometry (compare Supplementary Fig. 10b and Fig. 2).

**Fig. 4.**
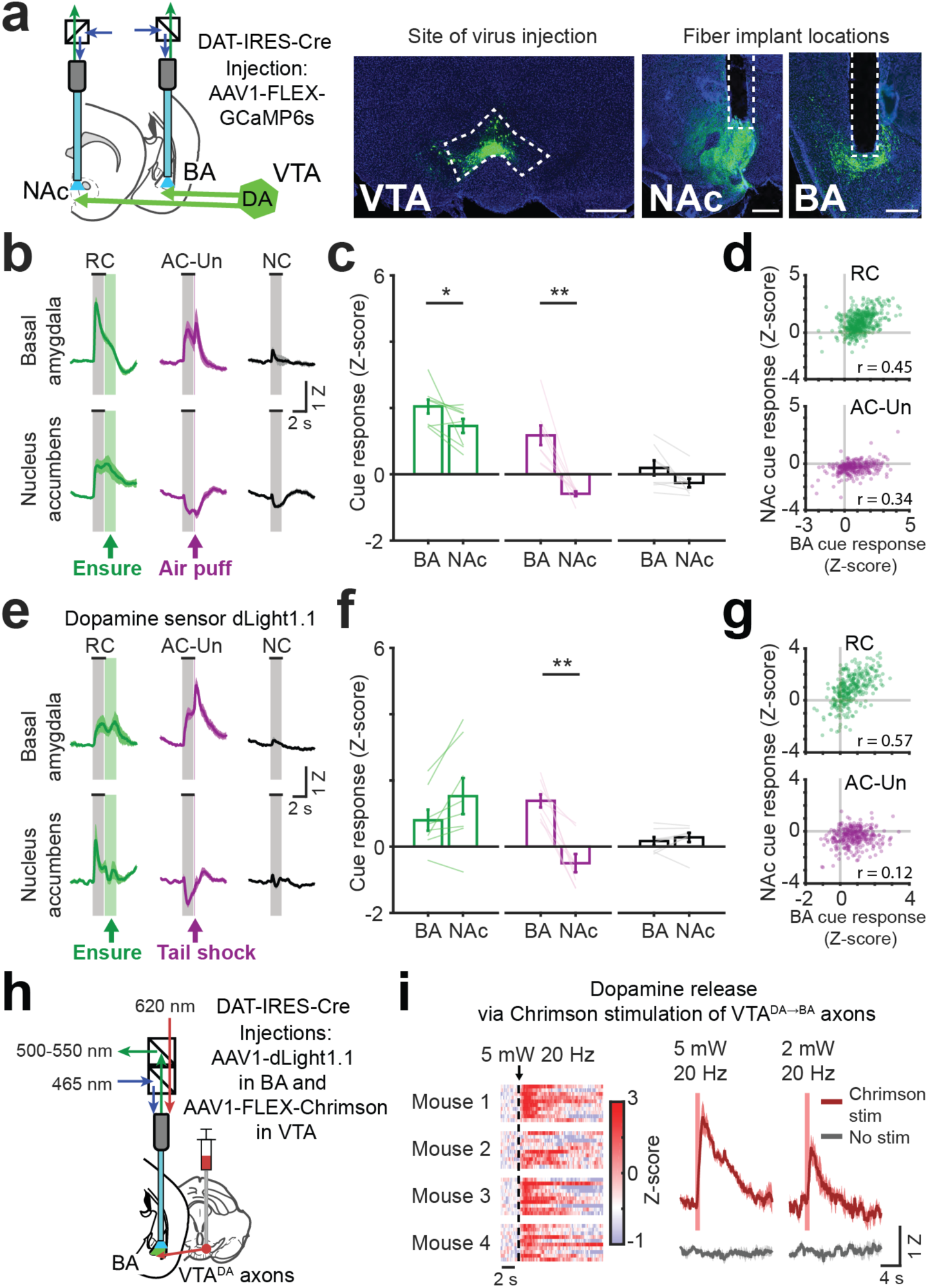
Opposite responses to aversive cues in simultaneous recordings of VTA dopamine axons or dopamine release in BA and NAc. **a,** *Left*: schematic of simultaneous fiber photometry recordings from VTA axons in nucleus accumbens (NAc) and in BA. *Right*: example histology showing VTA injection of AAV1-hSyn-FLEX-GCaMP6s and fiber placements above axon fields in NAc and BA. Scale bar = 0.5 mm. **b,** Mean cue responses of VTA axons in BA and in NAc. Mean ± s.e.m. across 8 sessions from 4 mice. **c,** Comparison of cue response magnitudes in BA and NAc (n = 4 mice, mean ± s.e.m across 8 sessions, * p < 0.05, ** p < 0.01, Wilcoxon sign-rank). **d,** Scatter plot of single-trial cue responses (averaged across the stimulus period) from simultaneous recordings of VTA axons in BA and in NAc, during presentations of the RC (*top;* Pearson’s r: 0.45, p < 0.0001, 500 trials from 4 mice) and the AC-Un predicting air puff (*bottom;* Pearson’s r: 0.34, p < 0.0001, 293 trials from 4 mice, 2 sessions per mouse). **e,** Mean cue responses from simultaneous recordings of the dopamine sensor dLight1.1 in BA and in NAc. Mean ± s.e.m. across 8 sessions from 4 mice. **f,** Comparison of cue response magnitudes of dopamine sensor in BA and NAc of hungry mice (mean ± s.e.m across 8 sessions from 4 mice, ** p < 0.01, Wilcoxon sign-rank). **g,** Scatter plot of single-trial cue-evoked dLight1.1 responses (averaged across the stimulus period) during presentations of the RC (*top;* Pearson’s r: 0.57, p < 0.0001, 326 trials from 4 mice) and the AC-Un predicting tail shock (*bottom;* Pearson’s r: 0.12, p < 0.0001, 320 trials from 4 mice, 2 sessions per mouse). **h,** Schematic of dLight1.1 fiber photometry recordings in BA and stimulation of VTA axons in BA using the red-shifted excitatory opsin, Chrimson. **i,** *Left:* heatmap showing single-trial dLight1.1 responses recorded in BA of four mice following Chrimson stimulation of VTA^DA→BA^ axons (1 s duration, 20 Hz, 5 mW, 620 nm). *Right:* mean dLight1.1 response in BA following stimulation at two light intensities (5 mW and 2 mW; 1 s duration, 20Hz; s.e.m. across 4 mice).

We continued to track the activity of VTA^DA→BA^ axons across daily sessions after replacing the passively avoidable quinine outcome with unavoidable airpuff, and then by unavoidable tail shock. We found that individual axons that were previously responsive only to the RC acquired responses to the AC-Un paired with air puff (Fig. 5b-d; Supplementary Fig. 10e,f; Supplementary Movie 1). AC-Un responses persisted in sated mice (Fig. 5b,e,f). These findings were even more pronounced when the AC-Un was paired with tail shock, now resulting in response preferences of individual VTA^DA→BA^ axons for the AC-Un (Fig. 5g,h). Across all mice, we observed a transition from response biases towards the RC when the task involved passively avoidable quinine, to unbiased or slightly AC-Un-biased responses when the task involved tail shock (Supplementary Fig. 10e). Furthermore, individual VTA^DA→BA^ axons became significantly more responsive to this AC-Un following satiation (Fig. 5i). Axons that responded preferentially to the AC-Un often showed weaker responses to both of the other cues (likely reflecting stimulus generalization), and these non-specific cue responses increased after satiation (Fig. 5g). However, a differential increase in AC-Un responses in the sated state persisted even after accounting for changes in overall cue responsivity across states (Supplementary Fig. 10f).

We were able to track many of the same axons across three weeks of training. The same axons were activated by the AC-Un paired with air puff and later with tail shock (Fig. 5j). Most VTA^DA→BA^ axons were activated by both the RC and AC-Un, while smaller subsets were selectively driven by either the RC or AC-Un (Fig. 5k,l). In almost all cases, VTA^DA→BA^ axons showed relatively weak responses to the NC (Fig. 5l). This sensitivity of individual VTA^DA→BA^ axons to appetitive and aversive cues predicts an increase in dopamine signaling in a given target BA neuron during presentation of any motivationally salient cue, regardless of valence.

### Excitatory neurons in BA also acquire responses to motivationally salient aversive cues

In Fig. 1, BA excitatory neurons were not preferentially responsive to stimuli lacking motivational salience, including the AC-Av and NC after training, and any visual stimuli prior to their association with salient outcomes. Given that, on average, VTA^DA→BA^ axons and BA excitatory neurons displayed similar insensitivity to non-salient cues, and that VTA^DA→BA^ axons were activated by the AC-Un but not by the AC-Av, we assessed whether certain excitatory BA neurons downstream of VTA^DA→BA^ axons would develop AC-Un response preferences^4^ (Fig. 6a). Following training on the task involving unavoidable aversive outcomes, a substantial number of BA neurons became preferentially active during the AC-Un (Fig. 6b-e; Supplementary Fig. 11a). As was the case with VTA^DA→BA^ axons, satiation induced a selective attenuation of RC responses but *not* of AC-Un responses across the population of BA excitatory neurons (Fig. 6f-h; Supplementary Fig. 11b-d), suggesting that satiation may promote a more defensive state reflected by a population-level shift in sensitivity towards aversive cues (Fig 6g; Supplementary Fig. 11e,f). These changes in response magnitude following satiation were not caused by decreased arousal, as analyses matched for pupil dilation or locomotion across states produced similar results (Supplementary Fig. 12; satiation did, however, decrease the overall proportion of cue-responsive BA neurons, Supplementary Fig. 11g).

Unlike individual VTA^DA→BA^ axons, but consistent with previous literature^4^, many BA neurons showed selective activation by either the RC or the AC-Un (Fig. 6c-e,i, Supplementary Fig. 11a). These neurons were often in close proximity (<50 μm apart), without any obvious fine-scale organization (Fig. 6j-k). This finding that distinct, intermingled subsets of neurons are activated by appetitive or aversive cues was true even when only considering a retrogradely labeled subset of NAc-projecting BA neurons (Supplementary Fig. 13a-h), consistent with the diversity observed in previous electrophysiology studies^37^. This intermingled arrangement of functionally distinct sets of BA excitatory neurons could allow both sets to receive salience signals from the same VTA^DA→BA^ axons during presentation of both appetitive and aversive cues.

### Comparisons of cell body recordings across subregions of BA

Previous studies suggested differences in valence processing across the anterior-posterior axis of BA^29^. We assayed for functional differences in aBA vs. pBA by separately analyzing data from GRIN lens implants in each of these subregions (Supplementary Fig. 1b and 14). When the RC was maximally salient during the hunger state, we observed spatial differences in RC responses: neurons exhibited larger RC responses in aBA vs. pBA and more frequently exhibited RC response preferences (Supplementary Fig. 14a,c,e). In contrast, when the AC-Un was maximally salient during satiety, neurons exhibited larger AC-Un response magnitudes in pBA vs. aBA and more frequently exhibited AC-Un response preferences (Supplementary Fig. 14b,d,f).

As certain prior studies^29^ comparing subregions of BA assessed activation by unconditioned stimuli (vs. predictive cues), we also asked whether aBA and pBA neurons respond differently to reward delivery vs. tail shock. The mean activity time courses of individual neurons in aBA and pBA showed a mixture of responses to cues and outcomes (Supplementary Fig. 15a). We therefore deconvolved the activity of individual BA neurons and constructed a generalized linear model^21^ to estimate the relative contribution of cues, unconditioned stimuli, and behavioral variables (see Methods and Supplementary Fig. 15b). We then identified neurons for which Ensure or tail shock delivery explained a significant amount of the activity (Supplementary Fig. 15b-d). We found that a larger proportion of neurons was activated by Ensure delivery in pBA vs. aBA (Supplementary Fig. 15e). In contrast (but consistent with analyses using a different method to identify cue-responsive neurons, Supplementary Fig. 14a), a smaller proportion of neurons was activated by the RC and a smaller proportion was suppressed by the RC, Ensure delivery, or licking in pBA vs. aBA (Supplementary Fig. 15 e,f). Together, these findings of relatively greater activation and lower suppression by Ensure in pBA are consistent with previous immediate early gene studies assessing unconditioned stimulus-evoked activity in pBA vs. aBA^29^, while the opposite spatial pattern of responses to the RC is consistent with that observed for VTA^DA→BA^ axons (Supplementary Fig. 9a-d).

### Glutamate co-release by VTA^DA^**^→^**^BA^ axons differentially targets inhibitory neurons

The above findings provide initial clues as to how VTA^DA→BA^ axons might impact BA cell bodies across learning and changes in hunger state. To gain additional insight into the actions of VTA^DA→BA^ axons, we performed *in vitro* patch-clamp recordings in BA neurons during optogenetic photostimulation of VTA^DA→BA^ axon terminals (Fig. 7a). Surprisingly, we observed fast inward currents that indicated activation of ionotropic glutamate receptors, confirmed by blockade using a glutamate receptor antagonist (Fig. 7b). This finding is consistent with an enriched expression of the vesicular glutamate transporter (*Vglut2*) in the more medial portion of VTA^33, 38^, where VTA^DA→BA^ cell bodies are predominantly located^33^.

We found that one-third of all recorded BA neurons had VTA^DA→BA^ axon-evoked monosynaptic glutamatergic currents (30/90 recorded neurons; Fig. 7c). We distinguished between fast-spiking inhibitory, other inhibitory, and excitatory neurons using a combination of electrophysiological properties and targeting of inhibitory neurons expressing tdTomato under control of the *Dlx* promoter (Fig. 7d; see Methods). Fast glutamatergic currents were significantly more common and of larger amplitude in fast-spiking interneurons than in excitatory neurons (Fig. 7e,f). In fact, these fast currents were often strong enough to trigger action potentials in fast-spiking interneurons (Fig. 7g), but not in excitatory neurons (not shown). Similar proportions of functional VTA^DA→BA^ glutamatergic synapses were found in recordings throughout the anterior-posterior axis of BA (Fig. 7h,i). Thus, VTA^DA→BA^ axons can release both glutamate and dopamine (Fig. 4i), and may shape network dynamics and functional plasticity in intermingled sets of BA excitatory neurons encoding cues predicting rewarding or aversive outcomes.

## Discussion

In this study, we used two-photon imaging and fiber photometry in head-fixed mice to examine the acquisition and motivational state-dependent expression of responses to learned cues in BA neurons and in the understudied set of VTA dopaminergic axonal inputs to BA. Initially arbitrary visual stimuli did not elicit responses in BA neurons. Once associated with salient outcomes, these cues could elicit either active approach behaviors (e.g. operant licking) or active avoidance behaviors (e.g. pre-emptive increase in eye closure or locomotion). Accordingly, intermingled sets of BA neurons developed preferential responses to either salient appetitive or aversive cues, but not to non-salient stimuli. We hypothesized that VTA^DA→BA^ axons may provide a teaching signal guiding the acquisition of salient cue responses in BA. Consistent with this prediction, VTA^DA→BA^ axons were activated by rewards and unavoidable punishments and, after training, by cues that predicted these salient outcomes. Furthermore, VTA^DA→BA^ axon responses to food cues decreased while responses to aversive cues increased following the transition from hunger to satiety, consistent with opposite changes in cue salience across these motivational contexts. Using GRIN-based two-photon calcium imaging of VTA^DA→BA^ axons, we found that individual VTA^DA→BA^ axons were activated by stimuli predicting reward and those predicting unavoidable punishment. As discussed below, these patterns of activation of VTA^DA→BA^ axons may create a window of plasticity for strengthening or weakening of visual inputs relaying information regarding salient appetitive or aversive outcomes to BA (e.g. from visual association cortex, thalamus, and lateral amygdala^20^). Our findings suggest that VTA^DA→BA^ neurons are a major driver of dopaminergic actions in BA during associative learning of stimuli paired with rewards or punishments^7–9, 12^.

### Attentional signaling in basal amygdala

BLA neurons may signal the associability of sensory stimuli when there is a significant change in the environment, resembling a reinforcement signal proposed by Pearce and Hall^28, 39^ in which reinforcement learning is enhanced by unexpected outcomes including rewards, punishments, and omissions of expected outcomes. While the activity of some BA neurons may align with the Pearce-Hall model in that they respond to rewards that are either larger or smaller than expected^40^, we did not find such activity in VTA^DA→BA^ axons. Instead, VTA^DA→BA^ axon activity either did not change or decreased when expected rewards or punishments were omitted (reflected by a sustained reduction in activity following the omitted outcome). Thus, any observed increases in BA neuron activity upon omission of rewards or punishments^40–42^ are unlikely to be mediated directly by VTA^DA→BA^ axons. Rather, we suggest that phasic responses of VTA^DA→BA^ axons may specifically promote learning of those cues that are associated with appetitive or aversive outcomes requiring active approach (e.g. licking) or avoidance (e.g. blinking or increased locomotion), thereby invigorating anticipatory motor responses.

### Prior studies of dopaminergic actions in lateral and basal amygdala

Our findings suggest that VTA^DA→BA^ neurons are a major contributor to the established role of dopaminergic actions in BA on associative learning^7–9, 12^ and argue against recent suggestions that VTA dopaminergic neurons do not play a role in aversive conditioning^15^. When considering the role of dopamine in BLA on associative learning, it is important to differentiate between sources and effects of dopamine in LA vs. BA. Our results (Fig. 2a) and others (Allen Brain Institute connectivity mapping, e.g. Experiment 301732962) suggest that VTA dopaminergic neurons send dense projections to BA, but not LA. Consistent with these anatomical findings, microdialysis measurements show basal dopamine release in BA, but not LA^43^. While VTA axons that innervate intercalated cells of the amygdala may gate LA activity, we suspect that the main actions of dopamine release from VTA axons on second messenger signals occur primarily within BA.

Our findings are consistent with recent studies indicating that VTA^DA→BA^ neurons originate in medial VTA^33^, where dopamine neurons display stronger responses to aversive outcomes^16^ and exhibit electrophysiological properties that differ from NAc-projecting dopamine neurons located in lateral VTA^16^. Our simultaneous photometry recordings from VTA dopamine axons in BA and NAc showed that these two subsets had aversive cue responses of opposing signs and weak trial-by-trial response correlations, suggesting distinct sources of input and/or distinct modulation at axon terminals. A larger fraction of medial VTA dopamine neurons co-express *Vglut2*^33, 38^. Accordingly, we find that VTA^DA→BA^ axons make monosynaptic glutamatergic connections with BA neurons, particularly onto fast-spiking interneurons.

### VTA^DA^**^→^**^BA^ inputs may support context-dependent learning of appropriate actions in response to salient cues

Why might VTA dopamine axons selectively innervate BA but not LA? Previous studies^44^ suggest that VTA dopaminergic neurons are particularly important for guiding learning in downstream circuits that link cues and associated outcomes with *active* goal-directed behaviors in a motivational state-dependent manner. BA is also implicated in such context-dependent behaviors and in cue-action-outcome learning, while LA appears more critical for action-independent passive learning of cue-outcome associations^45, 46^. Indeed, VTA^DA→BA^ axons showed motivational context-dependent increases in activity in response to cues that invigorated motor actions (licking to the RC in food-restricted mice but not in sated mice, and blinking or locomotion to unavoidable aversive cues), but not to neutral and passively avoidable aversive cues. We confirmed that these VTA^DA→BA^ axon responses were tightly locked to cue onset and not to onset of motor activity, suggesting that their role in cue-action-outcome learning is not due to direct regulation of same-trial motor activity. VTA^DA→BA^ axon responses and some BA neuron responses to unavoidable aversive cues were *enhanced* following the transition from hunger (a foraging state) to satiety (a defensive state), similar to previous studies showing that hunger suppresses neural and behavioral sensitivity to aversive cues and outcomes^47, 48^.

### Potential mechanisms for gating learning of motivationally salient sensory stimuli

An understanding of the role of VTA^DA→BA^ axons in guiding plasticity in BA should ideally include rapid effects of VTA^DA→BA^ axon signaling on network activity and dopaminergic effects on molecular signaling pathways that permit longer-term plasticity. While prior *in vitro* studies have observed direct actions of dopamine on the excitability of BLA excitatory neurons^49^, the predominant effects of dopamine are likely mediated by D1 receptor-dependent increases in cyclic adenosine monophosphate (cAMP) and activation of protein kinase A. Thus, the activation of the same VTA^DA→BA^ axons by both appetitive and aversive cues and outcomes may drive valence-independent increases in levels of plasticity-promoting cAMP in a given target BA neuron. If so, and if a given BA neuron receives diverse sensory inputs prior to learning, how is it that intermingled BA cells become selectively responsive to only appetitive *or* aversive cues in our task?

VTA^DA→BA^ axons may promote the selective acquisition of appetitive or aversive responses by driving enhanced competition *between* opposite-valence excitatory neurons. An enhancement in network inhibition via VTA^DA→BA^ activation of fast-spiking BA interneurons could ensure that a BA excitatory cell’s responses to stronger inputs are enhanced while responses weaker inputs are further suppressed, similar to hypotheses regarding the role of VTA dopaminergic projections to prefrontal cortex in increasing signal-to-noise ratio^50^. Other studies involving different tasks and degrees of training have also observed BA neurons with mixed responses to appetitive and aversive stimuli^51^. Further, neutral cues drove weaker yet more promiscuous responses in appetitive- or aversive-coding BA neurons, possibly due to weaker neutral cue-evoked lateral inhibition (Supplementary Fig. 11a). Such inhibition may occur, in part, due to the weaker activation of VTA^DA→BA^ axons during neutral cues, possibly providing a mechanism underlying generalization of cue responses. Consistent with these hypotheses, Esber and colleagues found that ablation of VTA dopamine neurons resulted in more promiscuous responses to salient reward cues across a larger fraction of BLA excitatory neurons^9^ and posited that VTA^DA→BLA^ inputs are essential for driving aspects of BLA activity related to attentional orienting towards salient or surprising stimuli, regardless of stimulus valence.

### Anatomical and genetic markers of valence coding in BA

Several models have been proposed for predetermining which BA neurons will be biased to positive or negative valence (see ^4^ for a detailed review), including hardwiring of inputs signaling valence-specific unconditioned stimuli^52, 53^, valence biases in projection-defined BA neurons^37^, anatomical organization of valence along the anterior-posterior axis^29^, and genetic markers of valence preference^29, 54, 55^. While our study was not initially designed to resolve this question, our use of two-photon calcium imaging in retrogradely labeled BA neurons and in GRIN lens implants in anterior BA (aBA) vs. posterior BA (pBA) afforded us the opportunity to consider whether our data might provide useful information.

We found that roughly equal proportions of NAc-projecting BA neurons responded to reward cues or aversive cues, consistent with heterogeneity previously observed in this sub-population^37^. We also examined whether BA neurons and/or dopaminergic inputs exhibited response biases in pBA vs. aBA. Using a generalized linear model to isolate responses to reward delivery, we confirmed previous immediate early gene analyses^29^ suggesting that reward outcome responses are somewhat more common in pBA. In contrast, reward cue response magnitudes were larger in aBA, both for cell bodies and for VTA^DA→BA^ axons. Despite these differences, most of the state-dependent response properties of BA neurons and VTA^DA→BA^ axons were qualitatively similar between pBA and aBA. While differences across studies may relate to the exact techniques employed (e.g. our use of viral labeling of projection neurons) and behavioral conditions involved, these data highlight the diversity of responses within projection-defined populations of BA neurons as well as across the anterior-posterior axis. In future, an intersection of anatomical location and genetic markers for defining both BA neurons and their projection targets may isolate more homogenous populations of valence-coding BA neurons.

### Potential limitations of our study

A large fraction of the sensory input to BA arrives via the LA. Our 500 μm diameter GRIN lens implant in BA damaged portions of the LA. This is more apparent in the anterior BA compared to the posterior BA, as the latter lies directly below the lateral ventricle. Nevertheless, the proportions of BA neurons that exhibited cue responses in our dataset (∼ 30%) are in line with previous reports using *in vivo* electrophysiological recordings^37^, and did not appear to vary with distance below the GRIN lens (range: 100 – 300 μm). In addition, the functional properties of VTA^DA→BA^ axons, whose dendrites reside in the midbrain, are unlikely to be strongly affected by our imaging procedures.

### Future directions and clinical significance

Our work provides a novel platform for high-resolution tracking of the activity of VTA^DA→BA^ axons and genetically-defined sets of target neurons across days and weeks in a mouse model during operant conditioning. In the future, this platform for chronic two-photon imaging of specific BA neuronal subtypes can be combined with local optogenetic and pharmacologic manipulations of dopaminergic inputs. These efforts should help elucidate the roles of VTA^DA→BA^ inputs in the acquisition of responses to motivationally salient cues, and in diseases involving overestimation of the salience of appetitive or aversive cues.

## Acknowledgements

We thank S.J. Lee, K. McGuire, J. Reggiani, A. Fratzl, Drs. M. Uchida, N. Uchida, J. Assad, L. Liang, Y. Livneh, J. Zaremba, S. Zhang, and other members of the Andermann lab for useful feedback. We also thank N. Pettit and Dr. Y. Aponte for helpful advice regarding GRIN lenses. We thank R. Jozanovic, E. Bamberg, F. Finkel, T. Pottala, I. Shurnayte, and L. Rupert for help with mouse training and histology. We thank Drs. J. Madara and H. Fenselau for assistance with brain slice electrophysiology. We thank Dr. L. Tian for providing dLight1.1 DNA plasmid, the Boston Children’s Hospital viral core for virus packaging and the GENIE Project, HHMI, for use of GCaMP6. Authors were supported by NIH F32 DK112589 and NIH 5T32NS007484-15 (A.L.), NIH T32 5T32DK007516 (A.U.S.), NIH New Innovator Award DP2 DK105570 and R01 DK109930, a McKnight Scholar Award, a Pew Scholar Award, a Smith Family Foundation Award, and grants from the Klarman Family Foundation, the American Federation for Aging Research, the Boston Nutrition and Obesity Research Center P30 DK046200, and a Harvard Brain Science Initiative Bipolar Disorder Seed Grant, supported by Kent and Liz Dauten (M.L.A.).

## Author contributions

A.L. and M.L.A. designed experiments and analyses and wrote the manuscript. A.L., O.A., H.K., and C.C. performed two-photon imaging experiments. A.L., C.C., V.D. and K.F. performed photometry experiments. A.L. performed brain slice electrophysiology experiments. A.L., O.A., C.C., V.F., and K.F. performed surgical procedures. A.L. analyzed two-photon imaging data with assistance from H.K. and A.U.S. A.L. analyzed photometry data, brain slice imaging, and electrophysiology data.

## Methods

All animal care and experimental procedures were approved by the Beth Israel Deaconess Medical Center Institutional Animal Care and Use Committee. Animals were singly housed on a 12-hour light/dark cycle with standard mouse chow and water provided *ad libitum* unless specified otherwise. *In vivo* experiments were performed on adult mice (12-24 weeks): Emx1-Cre;CaMKIIa-tTA;TITL-Ai941 (10 male mice were used for in vivo experiments), DAT-IRES-Cre2 (26 male and female mice were used for fiber photometry experiments; 4 male mice were used for axon imaging experiments; 10 male and female mice were used for in vitro slice experiments), and wild-type (C57Bl/6; 4 male mice for in vivo fiber photometry of dopamine sensor).

After surgical procedures, all mice in our experiments were singly housed. All experiments were performed during the light cycle (though chronic food restriction and food delivery during the task likely drives strong food entrainment). Experiments were typically performed on consecutive days or with one or two days in between imaging sessions. Mice included in the photometry and two-photon imaging experiments were between 12 and 24 weeks of age.

### Behavioral training

Initial behavioral training was performed as previously described3–5. After at least one week of recovery from surgery, mice were chronically food restricted to maintain body weight at 85% of free-feeding body weight by feeding ∼2.5 g chow per day (training and testing typically lasted 4 – 8 weeks). During testing, when mice performed 300 trials and consumed ∼ 3 – 5 mL of Ensure for satiation, mice were fed ∼ 1.5 g chow per day to maintained 85% body weight. Mice were initially habituated to head-fixation on a 3D printed running wheel, then trained to associate licking a lickspout with Ensure delivery by triggering Ensure delivery upon performance of a lick. Licks were detected using a capacitance-sensing lickspout (3D printed with conductive filament connected to a capacitance sensor, MPR121; Adafruit). All behavioral training was conducted using MonkeyLogic3,6.

Following shaping of licking behavior, mice were trained on an operant Go-NoGo visual discrimination task as previously described3. Mice learned to discriminate full-screen square-wave drifting gratings of different orientations (2 Hz; 0.04 cycles/degree; 80% contrast). The same three orientations (0° for reward cue; 270° for aversive cue; 135° for neutral cue) were used for all mice (previous studies from our lab have shown that the food cue bias observed in Fig. 1 for BA neurons is observed in upstream neurons in lateral amygdala and postrhinal cortex, and in downstream neurons in insular cortex, regardless of the initial grating orientations paired with food3–5). All visual stimuli were designed in MATLAB and presented in pseudorandom order on an LCD monitor positioned 20 cm from the mouse’s right eye. Stimuli were presented for 2 s, followed by a 2-s response window, and then a 6-s inter-trial interval. In the initial version of the task involving a passively avoidable aversive outcome (quinine delivery; e.g. Fig. 1-2), the first lick occurring during the response window triggered delivery of Ensure (∼ 5 μL) during reward cue trials and triggered delivery of quinine (∼ 5 μL; 0.1 mM) during aversive cue trials.

Following initial data collection using the task described above and previously3–5, the same mice were re-trained to associate the aversive cue (i.e. the 270° visual drifting grating) with an unavoidable air puff (50 ms duration; onset at 100 ms after termination of visual stimulus) delivered to the left eye (i.e. contralateral to the visual LCD monitor). A compressed air tank was used to generate the air puff and delivery was regulated by a solenoid (Clippard). A CCD camera was used to image the eye receiving the air puff delivery for determination of eye blinks/closures (either a Flea3 camera from PointGrey for photometry experiments or a Dalsa M1280 CCD camera for two-photon imaging experiments was used). For fiber photometry experiments, an infrared LED array (CMVision IRS48) provided illumination for imaging of the eye. For two-photon imaging experiments, infrared light emitted from the eye was sufficient to illuminate the eye. Mice showed anticipatory blinking behavior within or after one day of training, indicating that they learned the stimulus-outcome association rapidly.

Following 3 to 4 days of training on the task involving association between the aversive cue and an unavoidable air puff (2 sessions per day; ∼ 100 air puffs delivered per day), the same mice were re-trained to associate the aversive cue with an unavoidable tail shock delivery. Tail shocks (0.3 mA; 2 x 50 ms; 100 ms inter-shock-interval) were delivered via two electrode pads (Covidien; Series S) wrapped around the base of the tail. Current was delivered using a stimulus isolator (Iso-Flex; AMPI). Training on this new version of the task was then conducted for another 3 to 4 days. For all experiments, including those involving tail shock delivery, locomotion was monitored using a custom rotary encoder (two IR beam breaks detected the motion of tabs on the side of the printed wheel). Mice showed tail shock cue-evoked anticipatory increases in running speed within or after one day of training, indicating that they learned the stimulus-outcome association rapidly.

### Variations in task structure

For some experiments, variations in the behavioral task described above were used as detailed here. To assess whether decreases in VTA^DA→BA^ axon activity exist following omission of expected outcomes – Ensure, air puff, tail shock – we included “catch” trials (Supplementary Fig. 8). In the case of aversive outcomes, a random subset of 33% of all aversive cue trials were omission trials, and otherwise the task was as described above (3 cues, RC, AC, NC, presented with equal frequency). For reward omissions, mice previously trained on the main task were presented with 100 reward cue trials, and reward was omitted in 33% of these trials.

To assess the effects of sham satiation, a separate set of mice was trained with two operant cues: an Ensure-predicting cue and an avoidable tail shock-predicting aversive cue. Mice learned to perform licks to obtain Ensure, and to withhold licks to avoid delivery of tail shocks. Sham sating consisted of mice spending 1 hour with the lick spout accessible, but with no delivery of Ensure. Following recording of cue responses before and after sham satiation sessions, we returned mice to the home-cage where they had free access to chow. Once mice had returned to their normal free-feeding body weight, they were again presented with cues, and photometry recordings were collected to assess responses following caloric repletion.

### Virus injection surgical procedures

In the majority of experiments, AAV1-hSyn-FLEX-GCaMP6s (University of Pennsylvania Vector Core) was injected into the VTA of DAT-IRES-Cre mice (150 nl, Bregma: AP: −3.2 mm, DV: −4.5 mm, ML ± 0.3 mm). For VTA terminal stimulation experiments, AAV1-hSyn-FLEX-Chrimson-tdTomato^7^ (Addgene) was injected unilaterally into the VTA of DAT-IRES-Cre mice using the same volume and coordinates as above, and the dopamine sensor AAV1-hSyn-dLight1.1^8^ (plasmid from Dr. Lin Tian; virus packaged at Boston Children’s Hospital viral core) was injected in unilaterally in the BA ipsilateral to the VTA injection (150 nl, Bregma: AP: −1.8 mm, DV: −4.8 mm, ML +3.2 mm). For experiments testing home cage satiation, AAV5-hSyn-FLEX-axon-GCaMP6s (axon targeted GCaMP6s^9^; Addgene) was injected into the VTA of DAT-IRES-Cre mice using the same volume and coordinates as above. For experiments involving simultaneous ipsilateral fiber photometry recordings from dopaminergic terminals in basolateral amygdala (BA) and nucleus accumbens (NAc), only unilateral injections of virus were performed. In a subset of brain slice experiments, we labeled interneurons by injecting AAV8-hDLX-GqDREADD-tdTomato (plasmid from Addgene; virus packaged at Boston Children’s Hospital viral core), which expressed tdTomato under control of the *dlx* promoter^10^.

### Fiber implantation surgical procedure

Optic fibers with a metal ferrule (400 μm diameter core; multimode; NA 0.48; 5.0 mm length; Doric Fibers) were implanted over anterior (Bregma: AP: −1.2 mm; DV: −4.7 mm; ML: ± 3.2 mm) and posterior (Bregma: AP: −2.2 mm; DV: −4.7 mm; ML: ± 3.2 mm) BA on opposite hemispheres (hemispheres were counterbalanced across mice). For mice with optic fibers implanted over NAc and BA, the fibers were implanted over the medial shell of the NAc (Bregma: AP: 1.4 mm; DV: −4.5 mm; ML: 0.75 mm) and the BA (Bregma: AP: −1.6 mm; DV: −4.7 mm; ML: 3.2 mm). For VTA terminal stimulation experiments and for home cage satiation experiments, fibers were implanted in the middle of BA (Bregma: AP: −1.8 mm; DV: −4.7 mm; ML: 3.2 mm). The fibers and a custom-made titanium headpost were fixed to the skull using C&B Metabond (Parkell). Mice were given at least 2 weeks to recover before behavioral training.

### Fiber photometry recording

Fiber photometry recordings were performed using head-fixed mice that were free to run on a circular treadmill. Fiber optic cables (1 m long; 400 μm core; 0.48 NA; Doric Lenses) were coupled to implanted optic fibers with zirconia sleeves (Precision Fiber Products). Black heat shrink material was placed around the fiber coupling to prevent external sources of light (e.g. from the visual stimulus) from interfering with recordings. Excitation and emission light was passed through a GFP fluorescence minicube (FMC3_E(460-490)_F(500-550); Doric Lenses). Excitation light (∼ 100 μW) was provided by a 465 nm LED (Plexon LED and driver) which was modulated at either 217 Hz or 319 Hz using TTL output from two lock-in amplifiers (SR830; Stanford Instruments). Emission light was collected by a femtowatt photoreceiver (Newport 2151), demodulated using a lock-in amplifier (SR830; Stanford Instruments) and digitized at 1 kHz sample rate (PCIe-6321; National Instruments). Data acquisition was controlled using a custom script in MATLAB (MathWorks).

### GRIN lens implantation and related surgical procedures

For BA cell body imaging, mice were implanted with a singlet gradient index (GRIN) lens (GRINtech, NEM-050-25-10-860-S-1.5p; 0.5 mm diameter; 6.5 mm length; 250 μm focal distance on brain side at 860 nm (NA 0.5); 100 μm focal distance on air side (NA 0.5); non-coated). For axon imaging, mice were implanted with a doublet GRIN lens (GRINtech, NEM-050-25-10-860-DM; 0.5 mm diameter; 9.89 mm length; 250 μm focal distance on brain side at 860 nm (NA 0.47); 100 μm focal distance on air side (NA 0.19)). In a subset of mice, a polyimide guide cannula^11^ (Doric Lenses) was implanted and the GRIN lens was inserted each day of imaging and recovered after each imaging session (allowing for reuse across multiple mice). GRIN lens implantation coordinates for cell body imaging of BA neurons in transgenic mice (Emx1-Cre;CaMKIIa-TTA;TITL-Ai94) in anterior BA (aBA), medial BLA (mBA), or posterior BLA (pBA) (relative to Bregma): AP: −1.2 mm (aBA); −1.8 mm (mBA); −2.1 mm (pBA), ML: −3.2 mm, DV: −4.6-4.8 mm. In all mice where BA neurons were imaged, NAc-projecting BA neurons were retrogradely labeled by injecting 150 nL of AAVretro-hsyn-DIO-h2b-mCherry^12^ in NAc at (relative to Bregma): AP: 1.4 mm, ML: −0.75 mm, DV: −4.3 mm.

For VTA dopamine axon imaging in BA, VTA dopamine neurons were infected in DAT-IRES-Cre mice by injecting virus (150 nL, AAV1-hSyn-FLEX-GCaMP6s) at (relative to Bregma): AP: −3.2 mm, ML: −0.3 mm, DV: −4.3 mm), and a GRIN lens was implanted above BA at (relative to Bregma): AP: −1.8 mm, ML: −3.2 mm, DV: −4.6 mm.

GRIN lens implantations were performed as previously described13. Briefly, mice were maintained under anesthesia (1.5-2.0% isoflurane) and body temperature maintained using a heating pad in a stereotaxic apparatus (Kopf Instruments, Model 940). A beveled 25-gauge needle (0.51 mm diameter, Fisher) was attached to the stereotaxic holder and zeroed on Bregma. To give the lens a snug fit and reduce brain motion artifacts, the needle was inserted slowly to a depth of 0.1 mm higher than the final depth of the GRIN lens. After this needle was removed from the brain, the GRIN lens was held with a bulldog serrafine clamp (Fine Science Tools, Cat. No. 18050-28) with heat shrink on the tips (to improve grip and prevent damage) and zeroed at Bregma without touching the skull (to avoid debris covering the lens surface). The lens was then slowly inserted in the hole made by the needle and down to its final depth, then secured to the skull by applying Metabond (Parkell) or UV-curable glue (Loctite) around the lens. A titanium head plate was centered over the GRIN lens and fixed to the skull using Metabond. Afterwards, a 3D-printed plastic funnel was cemented onto the headplate, which allowed for a water reservoir for the Nikon 16x water immersion objective. Additionally, this funnel was used to secure a light shield between the head and the objective using Velcro, to prevent collection of stray light from the LCD monitor. After the completion of the GRIN lens surgery, the top of the GRIN lens was protected by a cut-off tip of an Eppendorf tube (Fisher) and secured using Kwik-Cast (WPI). The mice were allowed to recover from surgery for at least two weeks prior to any behavioral training.

### Two-photon imaging

Two-photon imaging was performed using a resonant-scanning two-photon microscope (Neurolabware) at 15.5 frames/second and 796 x 512 pixels/frame. Imaging was performed with a 16x 0.8 NA water-immersion objective (Nikon) for mice implanted with 1.5 pitch GRIN lens (GRINtech: NEM-050-25-10-860-S-1.5p; see above) or a 4x 0.28 NA air objective (Olympus) for mice implanted with 1.0 pitch 2.6x GRIN lens (GRINtech: NEM-050-25-10-860-DM; see above). Imaging fields of view were at a depth of 100-300 μm below the face of the GRIN lens. A Mai Tai DeepSee laser or InSight X3 laser (Spectra-Phsyics) was used. We imaged using an excitation wavelength of 960 nm for all calcium imaging of BA cell bodies and 940 nm for VTA axon imaging (using pre-chirp compensation for dispersion as much as possible). Laser power ranged between 40–60 mW at the front aperture of the objective (the power at the sample was substantially less because of partial transmission via the GRIN lens). GCaMP6 signals were collected using a green emission filter (510/84 nm; Semrock). For all BA neuron imaging, a prior injection of AAVretro-H2b-mCherry in NAc allowed red labeling of nuclei of NAc-projecting BA neurons (red emission filter: 607/70 nm; Semrock). We collected both green and red light using PMTs (H10770B-40; Hamamatsu).

### Acute brain slice preparation

For slicing, a choline cutting solution (CCS) was used^14^: 93 mM Choline-Cl, 2.5 mM KCl, 1.25 mM NaH2PO4, 30 mM NaHCO3, 20 mM HEPES, 25 mM glucose, 2 mM thiourea, 5 mM Na-ascorbate, 3 mM Na-pyruvate, 0.5 mM CaCl2 and 10 mM MgSO4 (pH to 7.3–7.4). Afterwards, artificial cerebrospinal fluid (ACSF) was used during all recordings: 126 mM NaCl, 21.4 mM NaHCO3, 2.5 mM KCl, 1.2 mM NaH2PO4, 1.2 mM MgCl2, 2.4 mM CaCl2, 10 mM glucose. Mice were anesthetized (isoflurane inhalation), decapitated, and the brains rapidly removed and chilled in ice-cold CCS. Using a vibrating tissue slicer (Campden; 7000smz-2), acute coronal slices (275 µm thick) containing the amygdala were obtained. Slicing was performed in ice-cold CCS followed by immediate incubation in CCS at 37°C for 15 minutes and then further incubation in ACSF at 37°C for 30 minutes. Slices were then kept at room temperature in ACSF for 25 minutes to 5 hours before being used for recording. All solutions were continuously oxygenated with 95 O_2_ and 5 CO_2_.

### Slice recording conditions

Slices were continuously superfused (flow rate: 2-5 ml/min) with oxygenated ACSF. Neurons were visualized using an upright microscope (Axioskop 2 plus; Zeiss) equipped with infrared differential interference contrast (IR-DIC). All channelrhodopsin-assisted circuit mapping experiments were performed at room temperature.

### Brain slice electrophysiology

Channelrhodopsin was first expressed in dopamine neurons in VTA by injecting AAV5-hSyn-FLEX-ChR2 (150 nL; U. Penn. Vector Core) into DAT-IRES-Cre mice (8–10 weeks of age). Brain slices were obtained at least 4 weeks after virus injection. All electrophysiological recordings were collected and amplified via a Multiclamp 700B (Molecular Devices). Voltage-clamp recordings were low-pass filtered at 4 kHz and sampled at 10 kHz. Current clamp recordings were low-pass filtered at 8 kHz and sampled at 20 kHz. Signals were digitized using a Digidata 1321A (Molecular Devices) and acquired using pClamp 10 (Molecular Devices). For whole-cell recordings, the pipette internal solution consisted of the following (in mM): 140 K-gluconate, 10 NaCl, 10 HEPES, 1 MgCl2, and 0.1 EGTA, pH 7.3 (∼300 mOsm). Pipettes were pulled from borosilicate capillary glass (Warner Instruments) and had tip resistances of 2-5 MΩ when filled with internal solution. Whole-cell voltage-clamp recordings were performed with the membrane potential clamped at −70 mV. Stimulation of channelrhodopsin in presynaptic terminals was achieved using a 10 ms blue light pulse (470 nm; ThorLabs LED) controlled by a programmable pulse stimulator (Master-8; A.M.P.I.). Recordings were collected as individual 10-second-long “sweeps’’ (typically 10–20 sweeps per cell) during which 5 light pulses were given every 500 ms (2 Hz). In some cases, 30 seconds were allowed between sweeps to allow for recovery of depression of currents (particularly in excitatory BA neurons). At the end of the recording, the firing properties of the neuron were obtained by switching to current-clamp configuration and injecting depolarizing steps of current to evoke action potentials. To identify whether synaptic inputs were monosynaptic^15^, in a subset of experiments, we recorded synaptic currents following application of tetrodotoxin (1 μM; TTX) to block action potentials and then additional application of 4-aminopyridine (100 μM; 4-AP) to depolarized terminals and restore monosynaptic evoked currents.

### Brain tissue preparation and immunohistochemistry

For all experiments that involved stereotactic injections of virus or implantation of a fiber or GRIN lens, we verified infection in the desired brain region and correct implantation of fibers and lenses above the desired brain region. Preparation of brain tissue for histological analysis is detailed below.

Mice were terminally anesthetized with tribromoethanol (Sigma Aldrich) and transcardially perfused with phosphate-buffered saline (PBS) followed by 10% neutral buffered formalin (Fisher Scientific). Brains were extracted, cryoprotected in 20% sucrose, and sectioned coronally on a freezing sliding microtome (Leica Biosystems) at 40 μm. Brain sections were washed in 0.1 M phosphate-buffered saline pH 7.4, blocked in 3% normal donkey serum/0.25% Triton X-100 in PBS for 1 hour at room temperature and then incubated overnight at room temperature in blocking solution containing primary antiserum. Afterwards, sections were washed in PBS and incubated in Alexa fluorophore-conjugated secondary antibody (Molecular Probes, 1:1000) for 2 h at room temperature. After several washes in PBS, sections were mounted onto gelatin-coated slides and fluorescent images were captured with an Olympus VS120 slide scanner microscope. We used the following primary antibodies: rabbit anti-dsRed (1:1000, Clonetech, 632496), rat anti-mCherry (1:1000, ThermoFisher, M11217) and chicken anti-GFP (1:1000, Life Technologies, A10262).

### Data analysis

All data analyses were performed using custom scripts in MATLAB (Mathworks) and ImageJ (NIH).

### Image registration and timecourse extraction for two-photon imaging experiments

Image registration and extraction of regions of interest (ROIs) for in vivo two-photon calcium imaging of BA cell bodies was performed using a previously published software pipeline written in MATLAB3,4. Briefly, we first downsampled images spatially by a factor of two. The red channel containing the signal from nuclear-labeled NAc-projecting neurons was used for image registration as it provides a sparse and stable fluorescent signal across time. To correct for x-y motion, each frame from a 30-minute imaging run was registered to a reference image (average of 1000 frames within a run) using efficient subpixel registration methods16. Within each imaging session, the reference image from each run (2-4 runs/session) was registered to the reference image of the first run of the day and was used to correct for across-run x-y shifts. VTA axon imaging movies were also registered as described above, but the green channel was used for registration as no nuclear labeling was used for these experiments.

We used PCA/ICA to extract masks of pixels with correlated activity, corresponding to individual axons or cell bodies^17^. By default, we used only the top 75% of pixels^18^, but we screened each prospective ROI and could edit the size of the mask, selectively removing the lowest probability pixels. Pixels found in more than one mask were excluded from further analyses. Timecourses were extracted by averaging each of the pixels within each binarized mask. We calculated neuropil activity as the median value of an annulus surrounding each ROI (inner radius: 15 pixels; outer radius: 50 pixels; pixels belonging to any other ROI were excluded from these annulus masks). This timecourse of neuropil activity was then subtracted from the activity timecourse of the associated ROI to create a fluorescence timecourse, F(t), where t is time of each imaging frame. The change in fluorescence was calculated by subtracting a running estimate of baseline fluorescence (F0(t)) from F(t), then dividing by F0(t): ΔF/F(t) = (F(t) - F0(t))/ F0(t), where F0(t) is a running estimate of baseline fluorescence calculated as the 10^th^ percentile of F(t) in the previous 32-second sliding window^19^. We then converted this ΔF/F(t) timecourse into a z-scored timecourse to allow for comparison of cell responses across states and mice. The z-scored timecourse of a cell was calculated by subtracting the mean of all timepoints (across all runs including both hungry and sated) and then dividing by the standard deviation of all timepoints. For visualization purposes, all example cue-evoked timecourses were re-zeroed by subtracting the mean activity in the 1 s prior to visual stimulus onset.

### Criteria for cue responsive neurons and analysis of cue response

To determine if a cell or axon was responsive to each cue, we used previously established, conservative criteria3–5, which are described in detail here. We calculated the cue-evoked response up to 65 ms (1 frame at 15.5 Hz) prior to the first lick detected (to avoid ascribing significant visual responses to cells or axons that were only responsive following motor actions). We performed a Wilcoxon sign-rank test for each frame post-stimulus onset against the 1-s baseline period prior to stimulus onset, with Bonferroni correction for multiple comparisons across frames (p < 0.01). If three consecutive frames were significantly different than the baseline period, a cell or axon was considered responsive to that cue. For all cells and axons with significant responses to at least one cue, a cell or axon’s preferred cue was determined as the cue evoking the largest response during the cue period. For estimation of a cell’s (or axon’s) mean cue-evoked response magnitude, and for estimation of a cell’s (or axon’s) response bias to a given cue, we averaged all trials containing presentations of that cue during the run to obtain a mean timecourse for that cell (or axon), and then the maximum response during the 2-s duration of the cue presentation was used as that cell’s (or axon’s) response magnitude. For estimation of the response bias index (Fig. 5k, 6i), we included cells (or axons) that had a significant response to at least one cue and set all negative values to zero.

### Analysis of distances between cells of the same functional type or of different types

We obtained the center of mass (centroid) from each ROI corresponding to an individual cell body. For all pairs of cells in a given category, we plotted the cumulative distribution function of pair-wise Euclidean distances between the pairs of x-y centroid coordinates.

### Using a generalized linear model (GLM) to identify components of cell responses

A Poisson GLM was fit to deconvolved activity of each cell, accounting for task and behavioral variables4,20 using the glmnet package implemented in Matlab. The deconvolved cell activity was first downsampled by a factor of two and then convolved with a Gaussian kernel (width 260 ms). A series of basis functions (Gaussian curves separated by 260 ms (i.e. 4 frames), full-width at half maximum = 260 ms) was made to span each task and behavioral variable. Task variable basis functions were generated for each cue type (reward cue, neutral cue, and unavoidable-tail shock aversive cue), further separated by trial type (hit, miss, correct reject, false alarm), and tiled across the entire cue duration (0-2 s). Behavioral variable basis functions were generated for times relative to (i) any stimulus offset (0 to 1 s from cue offset, reflecting general stimulus offset-related responses), (ii) reward or tail shock delivery (0 to 1 s), (iii) onset of a lick bout (0 to 1 s from bout onset, with lick bouts separated by ≥ 2 s), (iv) all other individual licks (one kernel at the time of each lick), (v) brain motion (the kernel convolved with the analog vector of x-y shifts determine during registration of movies) and (vi) locomotion (the kernel convolved with the running speed).

The GLM was fit on two-thirds of the data for each cell, using elastic net regularization (α = 0.01). We then used the GLM coefficients to measure the deviance explained on the remaining one-third of the data. Cells were determined to have activity that was locked to a given variable if (i) the deviance explained by the model was greater than 0.01 (note that the deviance explained is limited by the small number of behavioral variables) and (ii) the category of basis sets (i.e. food-cue, neutral-cue, aversive-cue, reward, tail shock, or licking basis sets) made up at least 5% of that explanatory value.

### Fiber photometry analysis

Photometry signals were sampled at 1 kHz, low-pass filtered below 100 Hz, and downsampled to 50 Hz. We calculated ΔF/F = (F − F0)/F0, where F0 is a running estimate of baseline fluorescence calculated as the 10th percentile of F(t) in the previous 30-second sliding window. The ΔF/F timecourse was z-scored by subtracting the mean of all timepoints (across all hungry and sated runs in a given session) and then dividing by the standard deviation of all timepoints. Signals for individual trials were then renormalized by subtracting the average of a 2-s baseline prior to each visual stimulus onset. For analysis of the responses to an individual cue, all trials containing presentations of that cue during a run were averaged to obtain a mean timecourse, and then the peak response during the 2-s duration of cue presentation was obtained.

To estimate response magnitudes to outcomes (Ensure and tail shock deliveries) from the fiber photometry, we fit monoexponential functions to the average cue timecourse (fitting using beginning at the peak of the cue response and extending until the end of the cue period) for each mouse individually. For each mouse, we then subtracted this fit from the photometry signal during subsequent post-cue timepoints to estimate the post-cue activity beyond that expected from residual cue response dynamics. The peak magnitude of this post-cue response (0-3 s post-cue offset) was then used to estimate responses to Ensure or tail shock.

For analyses of reward cue learning, we compared data from mice “During training” (performance is poor with high rate of false alarms), which refers to sessions during the first 3 days of training before introducing aversive trials, to data from mice that were “Trained” (performance is good with greater than 80% hit rate with less than 20% false alarms).

We note that our results are unlikely to be due to viral expression and changing levels of GCaMP over time because 1) we performed the experiments more than 6 weeks after initial infection (a period where increases in expression still occur but at a slower rate); 2) the transition from the task involving avoidable outcomes to the task involving unavoidable outcomes (and among tasks involving different unavoidable aversive outcomes) is fast -- several days -- and not likely explained simply by slow changes in expression (Supplementary Fig. 6); 3) we calculate fractional changes in fluorescence, which at least to some extent normalizes for changes in baseline fluorescence; and 4) most importantly, the fact that we observe consistent RC response magnitudes consistently across these days (see panel Supplementary Fig 6g) suggests that our findings are not due to changes in GCaMP6-related signal-to-noise across nearby sessions.

### Analysis of mouse behaviors

Eye closure during air puff experiments was determined from movies of the eye collected during photometry and two-photon imaging experiments21. The region of the video frames containing the eye was manually identified (at the beginning the movie when the eye was fully open) and then the average pixel intensity in this region was calculated for every frame. For fiber photometry experiments, closure of the eyelid resulted in increased signal intensity (reflected illumination of fur on eyelid by infrared LED is much brighter than the eye) which we used to determine eye blinks and persistent eye closure. Changes in eye closure during the 2-s visual cue or after air puff delivery were calculated relative to the 2-s baseline period prior to visual stimulus onset. Additionally, changes in baseline persistent eye closure across states (sated relative to hungry baseline) were also calculated to highlight the transition to a more defensive state in sated mice.

Locomotion was determined from the spherical treadmill rotary encoder. Changes in locomotion during the 2-s visual cue or after tail shock delivery were calculated relative to the 2-s baseline period prior to visual stimulus onset.

Pupil area was measured from the videography of the eye. A region around the eye was manually drawn and the center of the pupil was manually clicked for the first frame of the movie. The center and area of the pupil were then fit using a custom implementation of the starburst pupil detection from openEyes toolkit and a random sample consensus (ransac) algorithm. The center position from each previous frame was used to initialize the subsequent frame. In the case of videography of the eye acquired during two-photon imaging where the pupil was brighter than the rest of the eye (infrared laser light exiting out through the pupil), the starburst algorithm was set to use a decreasing gradient to fit the edge of the pupil. In the case of videography acquired during photometry, an infrared LED array illuminated the eye and the pupil was darker than the rest of the eye, the starburst algorithm was set to use an increasing gradient.

### Matching trials with similar pupil area or locomotion between states

To control for changes in arousal between hungry and sated runs, we determined average pupil area or locomotion in a 2-s window prior to visual stimulus onset5. For each trial during the hungry run, we attempted to find a trial in the sated run that had a mean pupil area that differed by less than 20%. These trials were deemed matching, removed from the set, and this process was repeated. All trials from the hungry and sated runs that were not successfully matched up were removed from further analyses. An identical procedure was performed to obtained trials that were matched for average pre-cue locomotion.

### Electrophysiology analysis

Individual sweeps from voltage-clamp recordings of light-evoked synaptic currents were normalized by subtracting the mean of the 2-s baseline period prior to light stimulus. Sweeps (10–20) were then averaged to obtain a mean timecourse. The peak amplitude of evoked synaptic current was used for determining if a cell received synaptic input from VTA axons (i.e. whether it was synaptically connected) and for comparison of synaptic current amplitudes between different cell types.

Neurons were classified as excitatory neurons, fast-spiking interneurons, or non-fast-spiking interneurons by morphology, membrane properties (resistance and capacitance), and action potential characteristics upon injection of depolarizing current steps in current-clamp configuration22. In addition, in a subset of experiments, we used slices from mice injected with AAV8-hDLX-GqDREADD-tdTomato (plasmid from Addgene; virus packaged at Boston Children’s Hospital viral core), which lead to expression of the red fluorescent protein, tdTomato, in the nuclei of inhibitory neurons10.

### Statistics

The numbers of samples in each group were based on those in previously published studies. Experiments were conducted by an investigator with knowledge of the animal genotype and treatment. As the behavior task was automated and trials were randomized, the investigator did not have prior knowledge of the timings of different trial types. Custom-written MATLAB analysis scripts allowed for data analysis in an automated and unbiased manner. All virus expression, optic fiber implants, and GRIN lens placements were verified by post hoc histology. Mice in which either the virus expression or optic fiber was not appropriately located were excluded from analysis. All data presented as bar and line graphs indicate mean ± s.e.m. with individual data points also plotted. Non-parametric statistical tests were used in the vast majority of cases. In a few cases with small sample sizes, parametric tests were used. Pair-wise comparisons were calculated using Wilcoxon rank-sum or Wilcoxon sign-rank tests, and multiple group data comparisons were calculated using Kruskal-Wallis tests with Bonferroni corrected post hoc comparisons between groups. Normality was not assessed as non-parametric tests were used. Statistical analyses were performed in MATLAB. Significance levels are indicated as follows unless otherwise specified: *p < 0.05; **p < 0.01; ***p < 0.001.

### Data and code availability

The data that support these findings and the code used for these analyses are available upon reasonable request.

---

**Supplementary Fig. 1.**
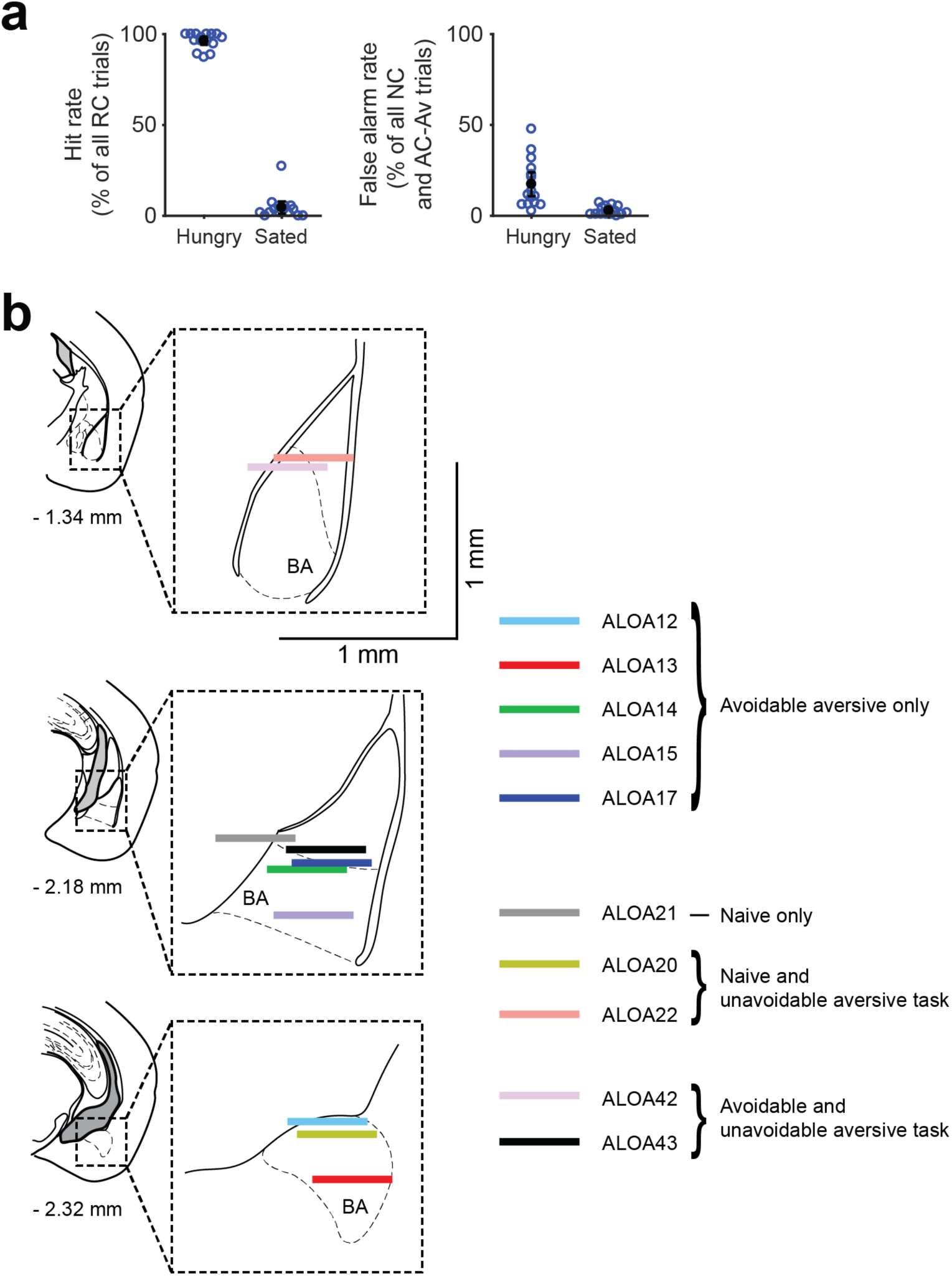
Go/NoGo behavior and GRIN lens locations. **a,** Behavioral performance of trained mice (n = 15 sessions from 7 mice) used for recordings in Figure 1. *Left:* percentage of correct responses across reward cue trials (‘hit rate’). *Right:* percentage of false alarms (incorrect licking after neutral or aversive cues) across all neutral and aversive cue trials. **b,** Confirmation of locations of the surface of the GRIN lens within the BLA (horizontal colored lines) for recordings related to Figures 1 and 6, determined from post-hoc histology (lenses are depicted at scale). The two-photon imaging fields of view were between 100 and 300 μm ventral to the lens surface. Coronal sections and coordinates relative to Bregma are based on Paxinos and Franklin’s mouse brain atlas (4^th^ edition).

**Supplementary Fig. 2.**
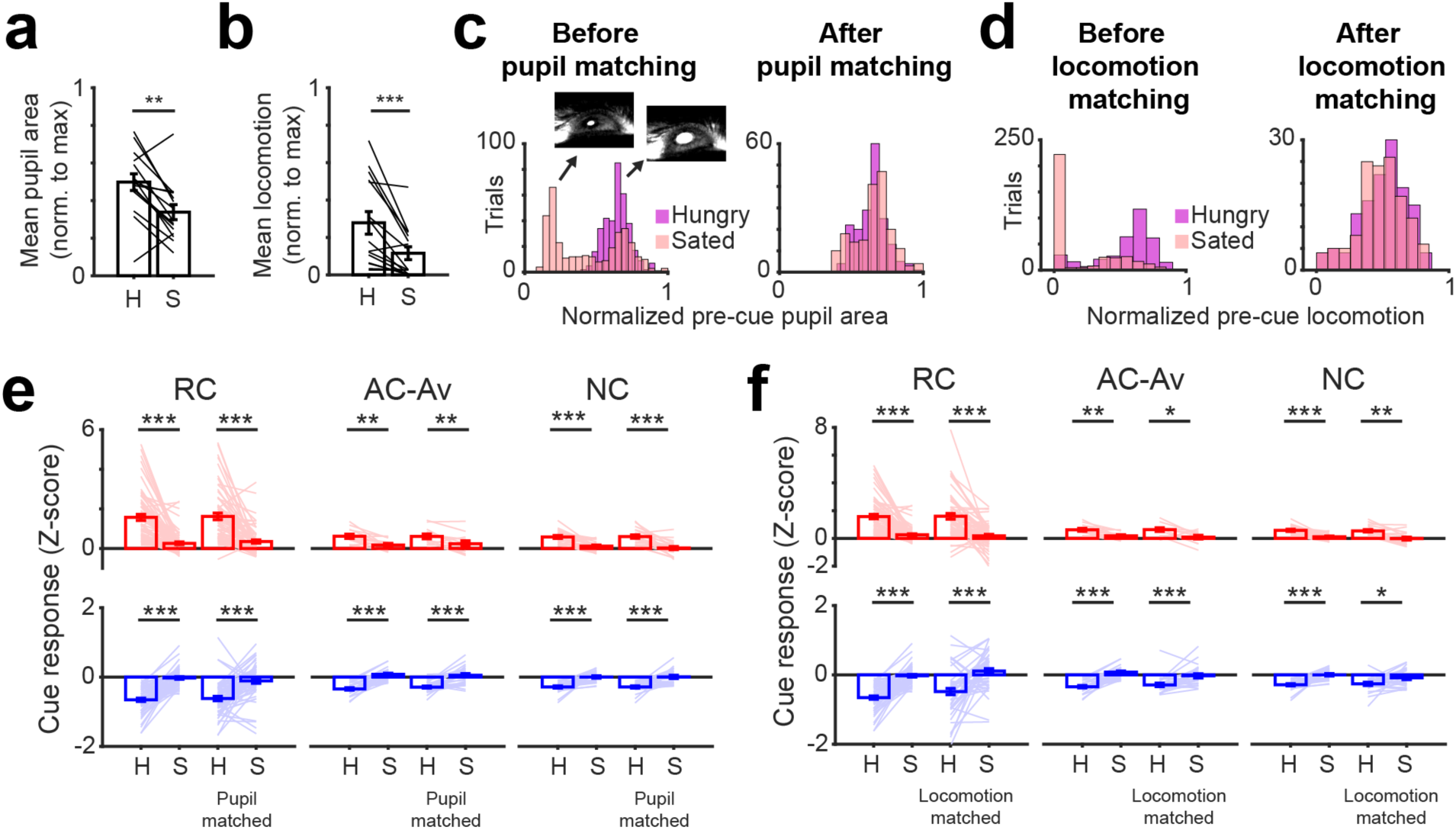
Changes in pupil area and locomotion across states do not account for observed changes in BA activity. **a,** Mean pupil area in the 2 s preceding a cue during hungry and sated states (normalized to maximum pupil area during the entire session across both states). Mean ± s.e.m. across 15 sessions from 7 mice. ** p < 0.01, Wilcoxon sign-rank. **b,** Mean speed of locomotion in the 2 s preceding a cue during hungry and sated states (normalized to maximum locomotion during the entire session across both states). Mean ± s.e.m. across 15 sessions from 7 mice. *** p < 0.001, Wilcoxon sign-rank. **c,** Histograms of pupil areas in the 2 s preceding onset of each cue during hungry and sated states from an example session before (*left*) and after (*right*) matching trials for pupil area (see Methods). Note that following matching of trials, distributions of pupil areas are now completely overlapping. Insets show moments of small (*left*) and large (*right*) pupil diameter (bright ellipse against a dark background, as infra-red laser light emitted from the pupil was used for tracking, see Methods). **d,** Histograms of relative locomotion speed in the 2 s preceding cues during hungry and sated states from an example session before (*left*) and after (*right*) matching trials for locomotion. **e,** Comparison of cue responses of BA neurons across hungry and sated states before and after matching for pupil areas. The finding of attenuated cue responses following satiation persisted even when matching pupil area distributions across states. ** p < 0.01, *** p < 0.001, Wilcoxon sign-rank. Z: Z-score. Mean response magnitudes were analyzed separately for neurons that were significantly activated (red) or suppressed (blue) by cue presentation. **f,** Comparison of cue responses of BA neurons across hungry and sated states before and after matching for locomotion. The finding of attenuated cue responses following satiation persisted even when matching locomotion distributions across states. * p < 0.05, ** p < 0.01, *** p < 0.001, Wilcoxon sign-rank. Z: z-score.

**Supplementary Fig. 3.**
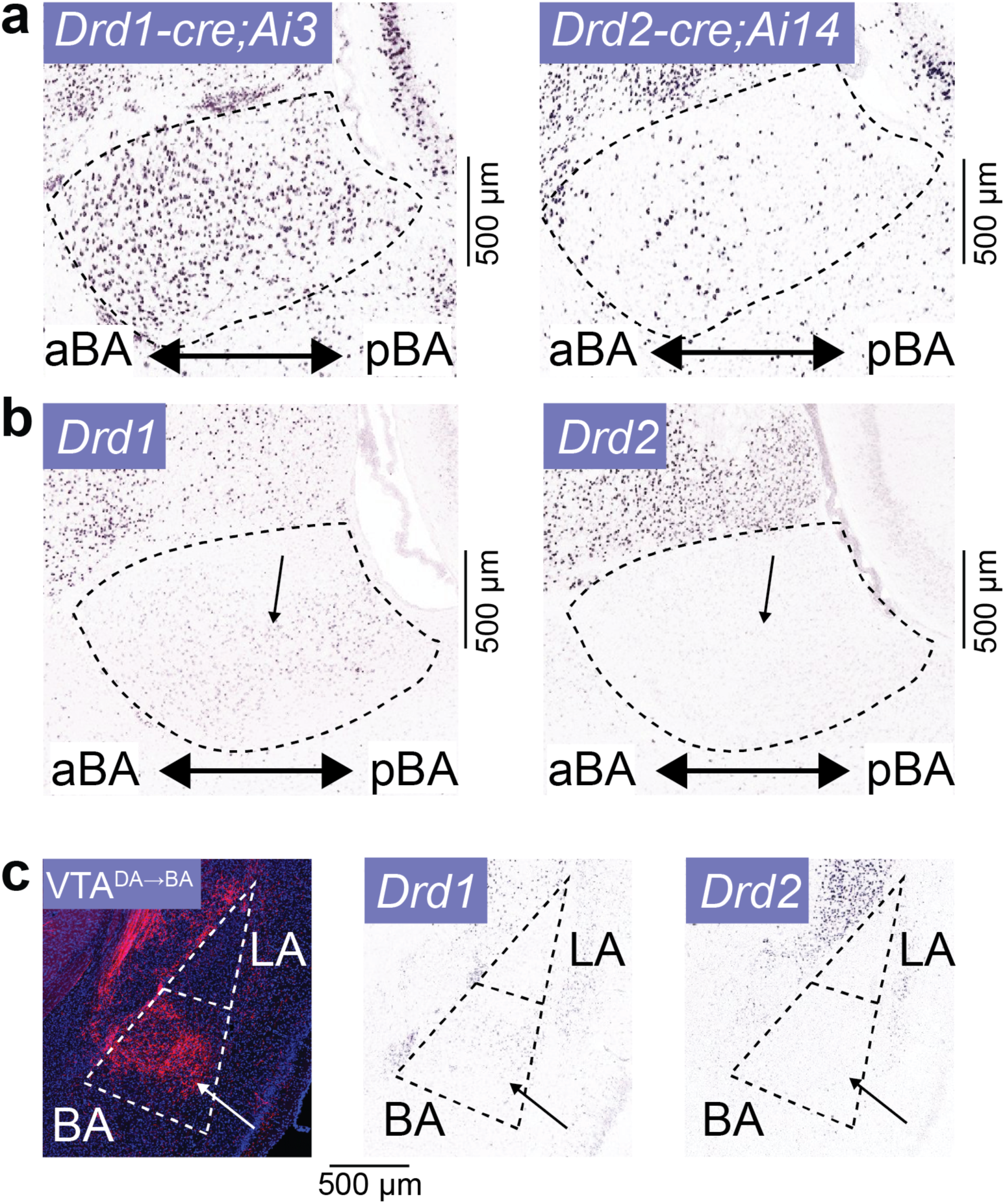
D1 receptor expression in basal amygdala. **a,** A cross of a red fluorescent reporter mouse with a D1 receptor Cre line (*left*) or a D2 receptor Cre line (*right*) provided initial evidence of dense D1 receptor mRNA expression and sparse D2 receptor mRNA expression across both anterior basal amygdala (aBA) and posterior basal amygdala (pBA) neurons, in sagittal sections (data modified from Allen Institute Brain Atlas; D1 receptor: image 3 [http://connectivity.brain-map.org/transgenic/experiment/81034295]; D2 receptor: image 1 [http://connectivity.brain-map.org/transgenic/experiment/100146223]). Because reporter fluorescence could reflect transient receptor expression during development, we also assessed mRNA expression levels in BA neurons in the Allen Brain Atlas (panel **b**). Scale bar = 0.5 mm. **b,** D1 receptor mRNA expression (*left*) and D2 receptor mRNA expression (*right*) in sagittal sections of BA. Note that expression of the D1 receptor was evident throughout anterior and posterior regions of BA, while D2 receptor expression was largely not detectable in BA (Allen Institute Brain Atlas; D1 receptor: image 3 [http://mouse.brain-map.org/experiment/show/71307280]; D2 receptor: image 3 [http://mouse.brain-map.org/experiment/show/81790728]). Scale bar = 0.5 mm. **c,** *Left:* mCherry-labeled VTA dopaminergic axons target BA but not lateral amygdala (LA) (for details, see Figure 2A). *Middle* and *right:* coronal sections demonstrating detectable expression of D1 receptor mRNA (*middle,* arrow) but not of D2 receptor mRNA (*right,* arrow) in BA (Allen Institute Brain Atlas; D1 receptor: image 24 [http://mouse.brain-map.org/experiment/show/352]; D2 receptor: image 26 [http://mouse.brain-map.org/experiment/show/357]). Scale bar = 0.5 mm.

**Supplementary Fig. 4.**
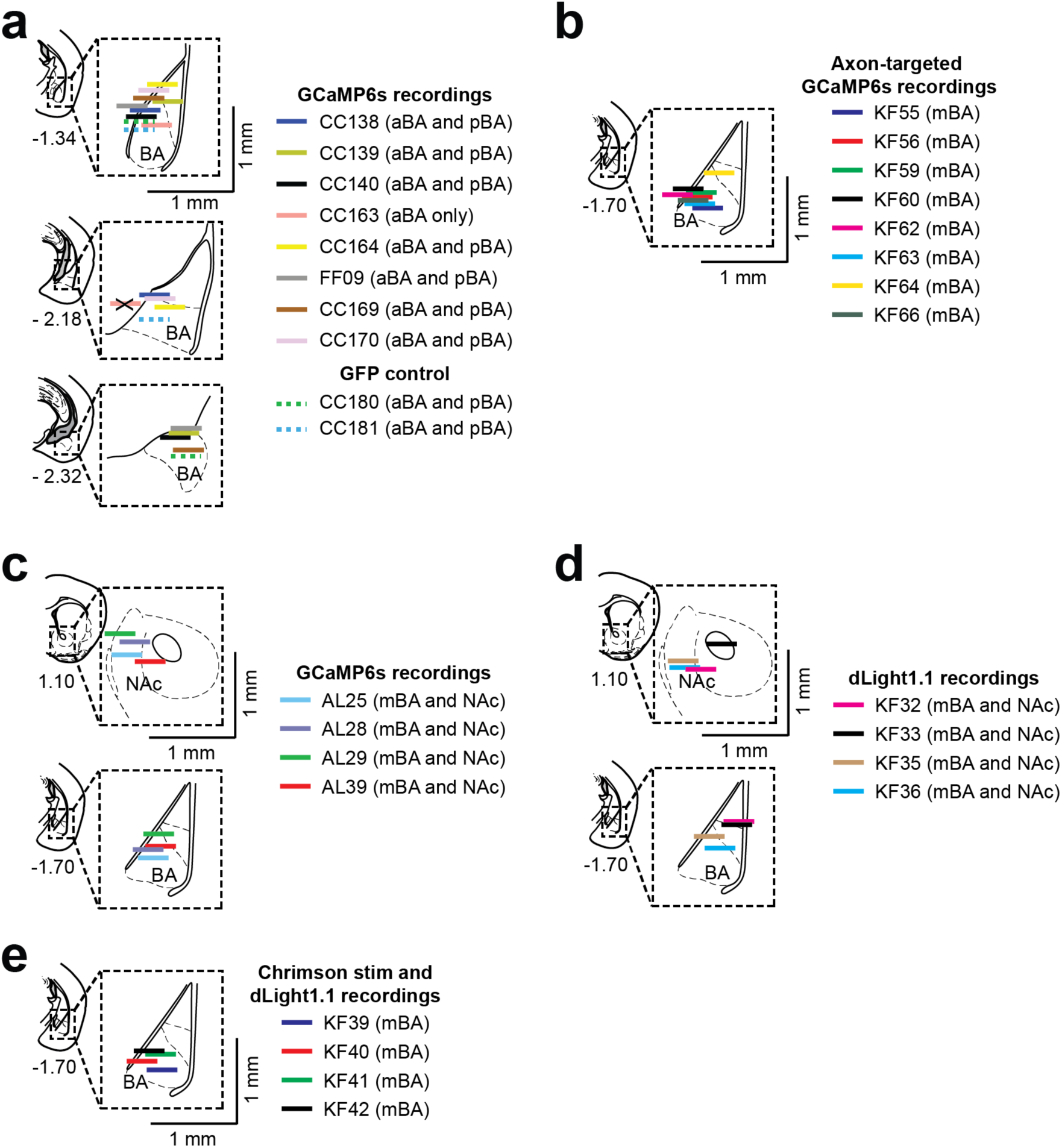
Implant locations for VTA^DA→BA^ fiber photometry experiments. **a,** Location of photometry fiber implants from all basal amygdala (BA) implanted GCaMP6s photometry mice (horizontal colored lines) determined from post-hoc histology (lenses are depicted at scale): 8 mice with bilateral BA implants targeted either to anterior BA (aBA) or posterior BA (pBA) regions, and mice used for two GFP control experiments. Photometry data from these mice are shown in Fig. 2c-e, Fig. 3c-I, Supplementary Figs. 5-9. Coronal sections and coordinates relative to Bregma are based on Paxinos and Franklin’s mouse brain atlas (4^th^ edition). ‘X’ marks a fiber implant excluded from subsequent analyses due to mis-targeting. **b,** Location of photometry fiber implants from all BA implanted axon-targeted GCaMP6s photometry mice for sham-satiation and home cage satiation experiments (horizontal colored lines) determined from post-hoc histology (lenses are depicted at scale): 8 mice. Photometry data from these mice are shown in Figs. 2f-h and 3j-l. Coronal sections and coordinates relative to Bregma are based on Paxinos and Franklin’s mouse brain atlas (4^th^ edition). **c,** Location of photometry fiber implants from all GCaMP6s photometry mice for comparing nucleus accumbens (NAc) vs. BA responses (horizontal colored lines) determined from post-hoc histology (lenses are depicted at scale): 4 mice with ipsilateral BA and NAc implants. Photometry data from these mice are shown in Fig. 4b-d. Coronal sections and coordinates relative to Bregma are based on Paxinos and Franklin’s mouse brain atlas (4^th^ edition). **d,** Location of photometry fiber implants from all dLight1.1 photometry mice for comparing NAc vs. BA responses (horizontal colored lines) determined from post-hoc histology (lenses are depicted at scale): 4 mice with ipsilateral BA and NAc implants. Photometry data from these mice are shown in Fig. 4e-g. Coronal sections and coordinates relative to Bregma are based on Paxinos and Franklin’s mouse brain atlas (4^th^ edition). **e,** Location of photometry fiber implants from all BA implanted dLight1.1 photometry mice for Chrimson optogenetic stimulation of VTA dopamine axon terminals (horizontal colored lines) determined from post-hoc histology (lenses are depicted at scale): 4 mice. Photometry data from these mice are shown in Fig. 4i. Coronal sections and coordinates relative to Bregma are based on Paxinos and Franklin’s mouse brain atlas (4^th^ edition).

**Supplementary Fig. 5.**
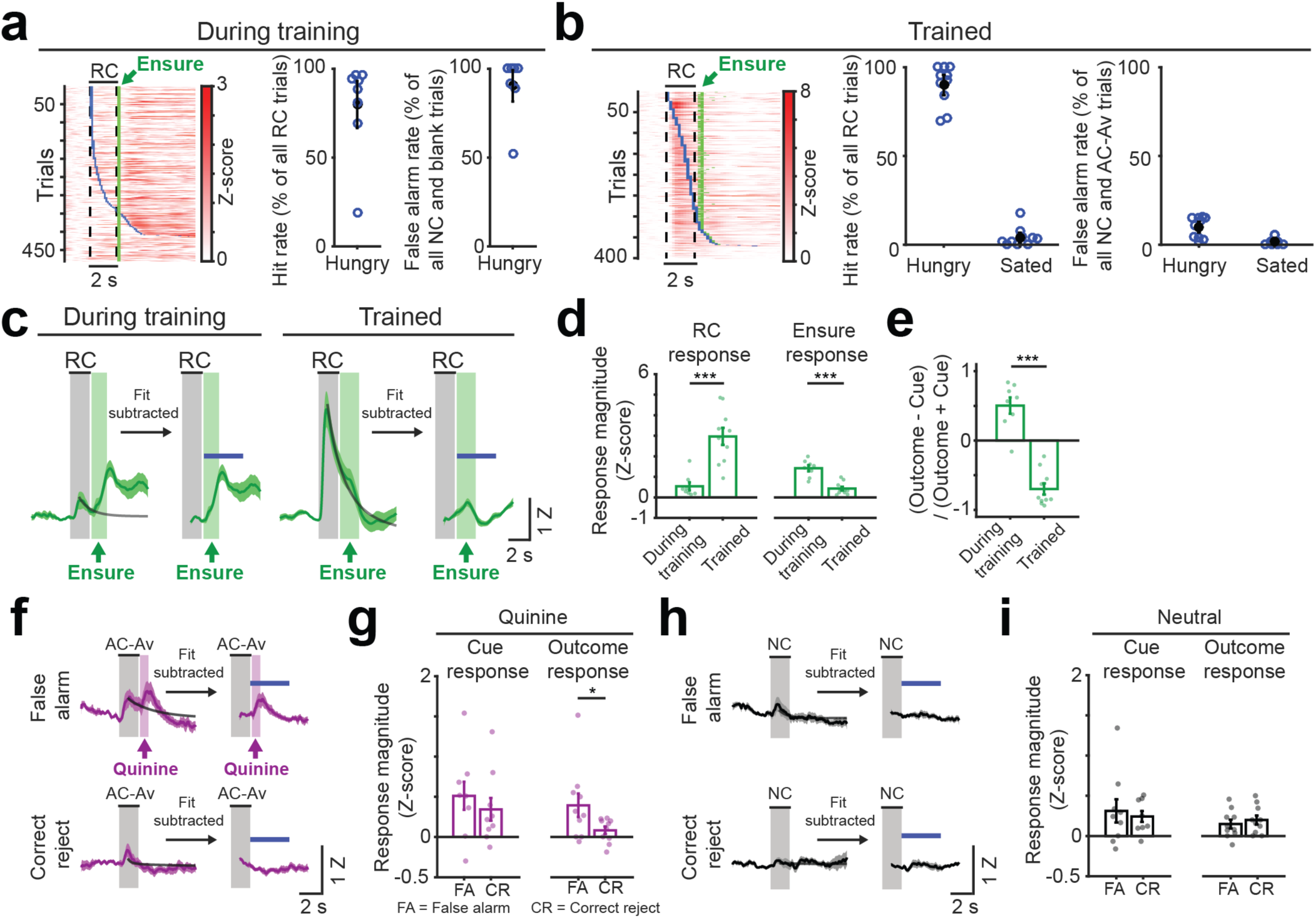
VTA^DA→BA^ axon responses during training and following completion of training. **a,** *Left*: all single-trial responses (rows) from 8 mice during Pavlovian training of RC paired with Ensure delivery (“During training” refers to sessions during the first 3 days of training before introducing aversive trials), sorted by onset of first lick following RC onset (blue ticks). Green ticks: time of Ensure delivery. Responses were mostly evident during Ensure consumption. This was especially pronounced when Ensure delivery was unexpected, as seen in rows with longer latencies to first lick. *Right*: behavioral performance of mice during training (n = 8 sessions, one session/mouse) used for recordings in panels **c-e**, below. Percentage of reward cue trials with licking during the cue (‘hit rate’) and percentage of false alarms (incorrect licking after neutral cues or during blank trials) were both high early in learning, indicative of poor discrimination between cues. **b,** *Left*: all single-trial responses (rows) from 10 mice following completion of training, sorted by onset of first lick following RC onset (blue ticks). Green ticks: time of Ensure delivery. Responses were locked to cue onset, not to motor response onset. *Right*: behavioral performance of trained mice (n = 10 sessions, one session/mouse) used for recordings in Figure 2d-e and in panels **c-e**, below. *Left:* percentage of correct responses across reward cue trials (‘hit rate’). *Right:* percentage of false alarms (incorrect licking after neutral or aversive cues) across all neutral or aversive trials. **c,** Mean response timecourses of VTA^DA→BA^ axons during training and following completion of training on task. Response timecourse of Ensure delivery related activity was obtained by subtracting a monoexponential fit of the cue response. The window used for analysis of Ensure delivery responses is indicated by a blue bar. Error bars: s.e.m. across mice. Z: Z-score. **d,** Comparison of cue responses and Ensure delivery responses during training vs. following training. *** p < 0.001, Wilcoxon rank-sum. **e,** The outcome response bias index shifted from positive to negative following training, indicating a shift in response from the time of Ensure delivery to the time of cue onset. *** p < 0.001, Wilcoxon rank-sum. **f,** Mean population response, averaged across those AC-Av trials early in training that involved licking during the behavioral response window (“false alarm”, thereby eliciting quinine delivery) or across trials without licking (“correct reject”, thereby eliciting no outcome). A monoexponential fit was subtracted to isolate the response magnitude to quinine delivery or no outcome. The window used for analysis of outcome responses is indicated by a blue bar. Error bars: s.e.m. across mice. Z: Z-score. **g,** Comparison of AC-Av cue and outcome responses for false alarm trials vs. correct reject trials, * p < 0.001, Wilcoxon sign-rank. **h,** Mean population response, averaged across those NC trials early in training that involved licking during the behavioral response window (“false alarm”) or across trials without licking (“correct reject”). In both cases, no outcome occurs following cue offset. A monoexponential fit was subtracted to isolate the response magnitude during the outcome window. The window used for analysis of outcome responses is indicated by a blue bar. Error bars: s.e.m. across mice. Z: Z-score. **i,** Comparison of NC cue and outcome responses during false alarm trials vs. correct reject trials.

**Supplementary Fig. 6.**
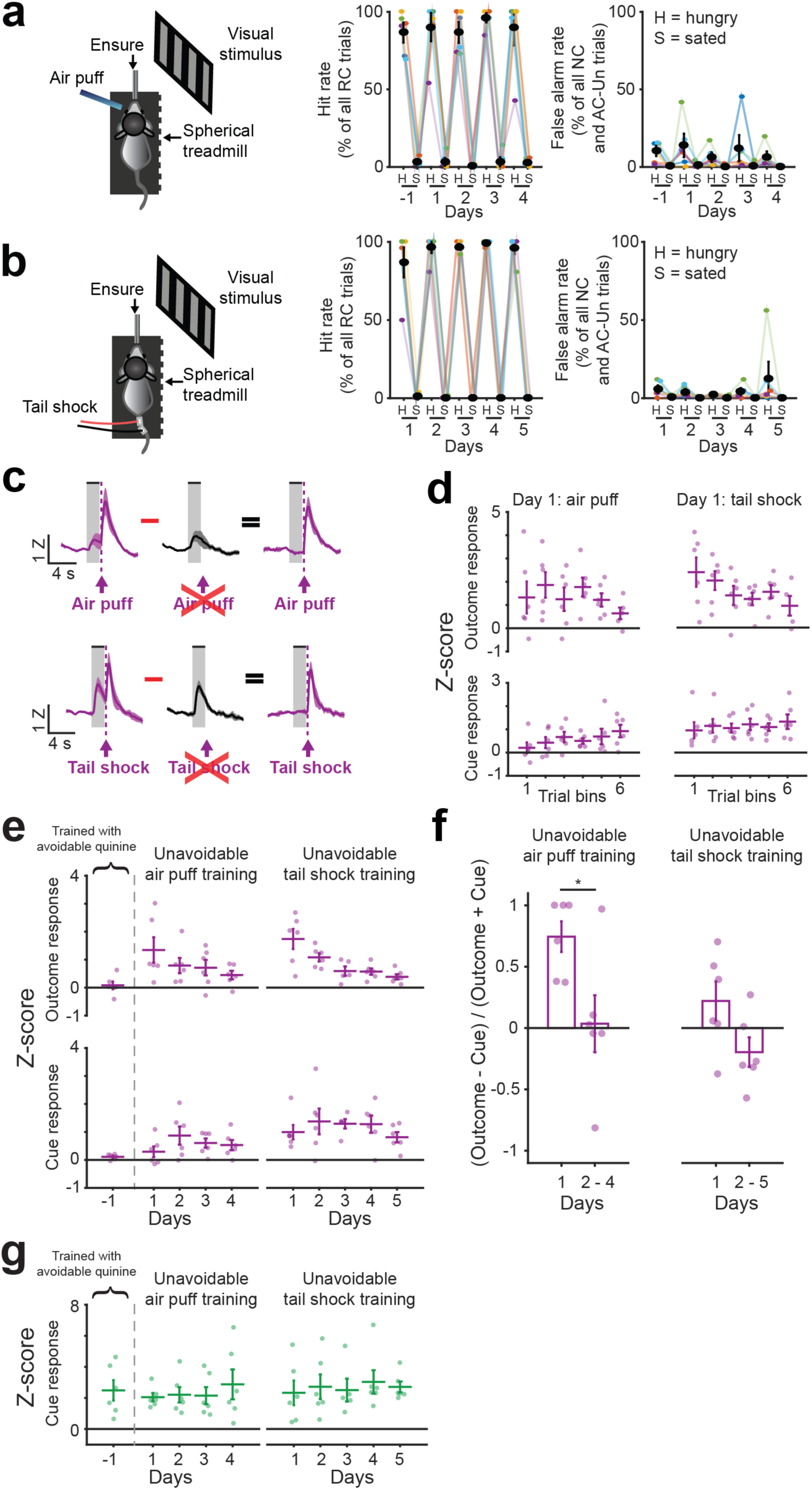
VTA^DA→BA^ axon responses during training with cues predicting air puff or tail shock. **a,** *Left:* schematic of delivery of air puff aversive outcome during the head-fixed Go/NoGo visual discrimination task. Behavioral performance of trained mice (n = 6 sessions, 1 session/mouse) remained high on the day preceding air puff training when mice were already trained on avoidable quinine task (day = −1) and during 4 days of training with air puff. *Middle:* percentage of correct responses across RC trials (‘hit rate’) during hungry and sated states. *Right:* percentage of false alarms (incorrect licking after neutral or aversive cues) across all NC or AC-Un trials during hungry and sated states. **b,** *Left:* schematic of delivery of tail shock aversive outcome during the head-fixed Go/NoGo visual discrimination task. Behavioral performance of trained mice (n = 6 sessions, 1 session/mouse) during 5 days of training with tail shock remained high. *Middle:* percentage of correct responses across RC trials (‘hit rate’) during hungry and sated states. *Right:* percentage of false alarms (incorrect licking after neutral or aversive cues) across all NC or AC-Un trials during hungry and sated states. **c,** Mean response timecourse on first sessions with aversive cue pairing with air puff (*top*) or tail shock (*bottom*) (s.e.m. across 6 mice). Trials in which we omitted the air puff (*top*) or tail shock (*bottom*) delivery did not have outcome responses and were subtracted from the trials with delivery of air puff (*top*) or tail shock (*bottom*) in order to isolate the response specific to the outcome. **d,** Outcome and cue response magnitudes on the first day of training with air puff delivery (*left*) or tail shock delivery (*right*) separated in 6 consecutive bins (∼ 10 trials per bin). Within the first day of training, outcome response magnitudes decreased whereas cue responses increased. Note that training with tail shock delivery followed training with air puff and therefore cue responses began at an already elevated magnitude. **e,** Outcome and cue response magnitudes across days of training with air puff (*left*) or tail shock (*right*). Across days, outcome responses continued to decrease while cue responses increased. **f,** An outcome bias was initially present on the first day of training with air puff delivery, but this bias showed a significant decrease following days of training. * p < 0.05, Wilcoxon sign-rank. **g,** Across all days of training with air puff (*left*) and tail shock (*right*) delivery, the average RC response magnitude remained stable.

**Supplementary Fig. 7.**
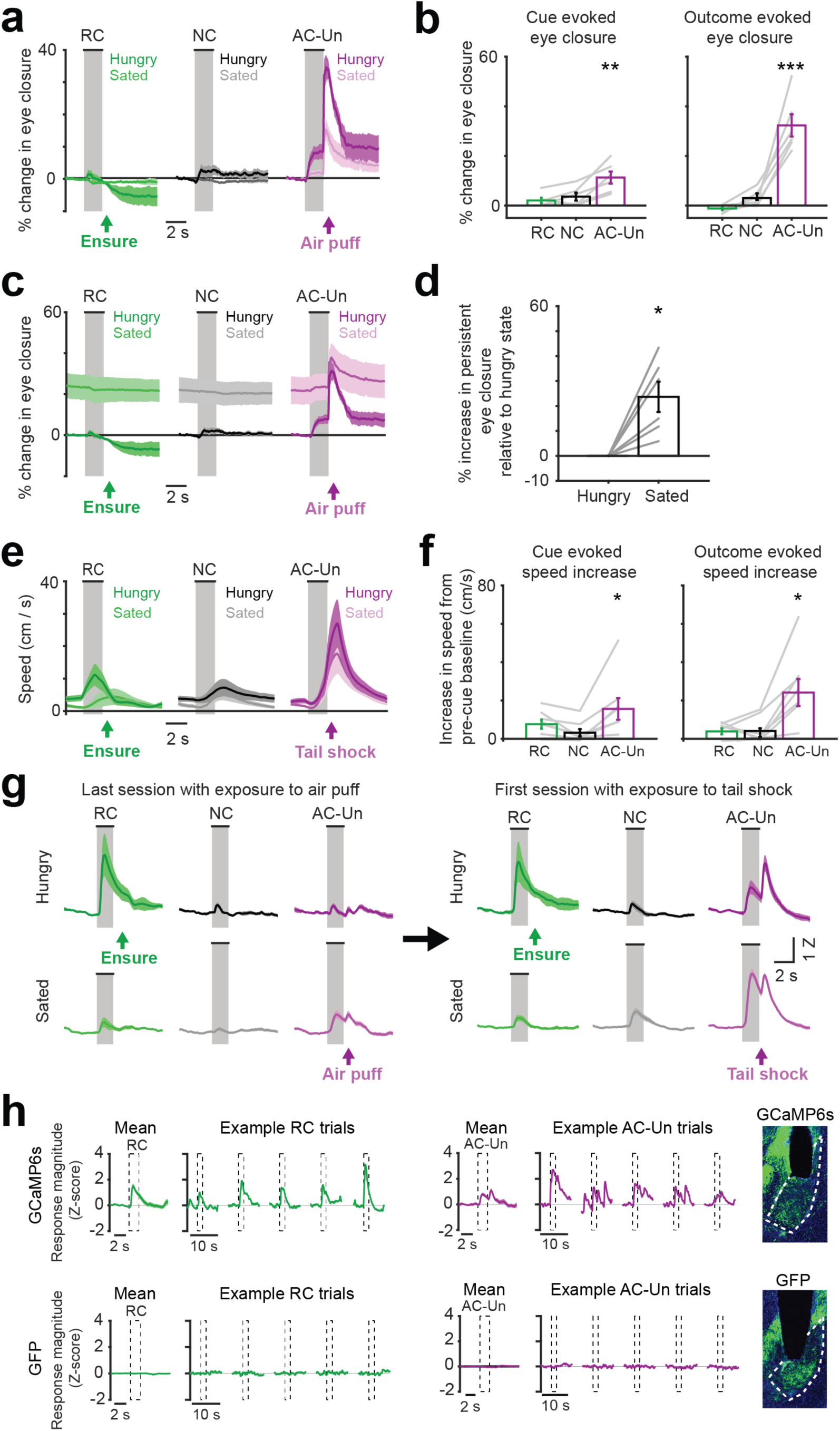
Behavior during tasks involving unavoidable aversive air puff or tail shock and related photometry controls. **a,** Percentage change in eye closure behavior for task involving unavoidable air puff, relative to pre-cue baseline period for reward cue (RC), neutral cue (NC), and aversive cue (AC-Un) trials. Error bars: s.e.m. across 6 mice. See Methods for details. **b,** *Left:* percentage change in eye closure during each cue period. *Right:* percentage change in eye closure after air puff delivery (n = 6, *** p < 0.001, ** p < 0.01, one-sample t-test against null distribution with mean of zero). Mean ± s.e.m. across 6 mice. **c,** Percentage change in eye closure for air puff experiments, relative to pre-cue baseline period from *hungry* trials. Note that persistent defensive eye closure increased with satiety. Note that VTA^DA→BA^ axon responses to unavoidable aversive outcome predicting cue (AC-Un) did not decrease with satiety, indicating that vision was not impaired in the contralateral eye receiving visual stimuli. **d,** Percentage increase in pre-cue persistent eye closure during sated trials relative to hungry trials. (n = 6, * p < 0.05, one-sample t-test again null distribution with mean of zero) **e,** Mean running behavior for task involving unavoidable tail shock during second session of training for reward cue (RC), neutral cue (NC), and aversive cue (AC-Un) trials. Error bars: s.e.m. across 8 mice (1 session/mouse). **f,** *Left:* change in speed during cue period relative to pre-cue baseline. *Right:* change in speed during tail shock delivery relative to pre-cue baseline (n = 8 mice, * p < 0.05, one-sample t-test again null distribution with mean of zero). Mean ± s.e.m. across 6 mice. **g,** *Left:* mean cue responses in fiber photometry recordings from VTA^DA→BA^ axons (4^th^ and final day of air puff exposure) in hungry mice and sated mice. Error bars: s.e.m. across 6 mice. Note that response to AC-Un progressively decreased across days (see also Supplementary Fig. 6; we speculate that this may reflect a decrease in motivational salience of the cue as mice develop learned helplessness), but continues to increase in sated sessions. *Right:* mean VTA^DA→BA^ cue responses (first session of tail shock exposure) in hungry mice. Error bars: s.e.m. across 6 mice. Z: Z-score. Note that the same mice previously learned the AC-Un association with unavoidable air puff and that the AC-Un response is rapidly restored upon exposure to the same visual cue now predicting unavoidable tail shock. **h,** *Top:* example GCaMP6s fiber photometry recording from VTA^DA→BA^ axons showing mean response and single-trial responses to reward cue (RC; *left*) and to cues predicting unavoidable air puff delivery (AC-Un; *right*). Note that Ensure reward delivery or air puff delivery occur following the second vertical dashed line. *Bottom:* mean and single-trial photometry traces from GFP controls do not show the same transient events observed in GCaMP6s recordings. Images at right: example histological reconstructions of fiber placements over BA and fluorescence of VTA^DA→BA^ axons for GCaMP6s (*top)* and GFP (*bottom)* experiments.

**Supplementary Fig. 8.**
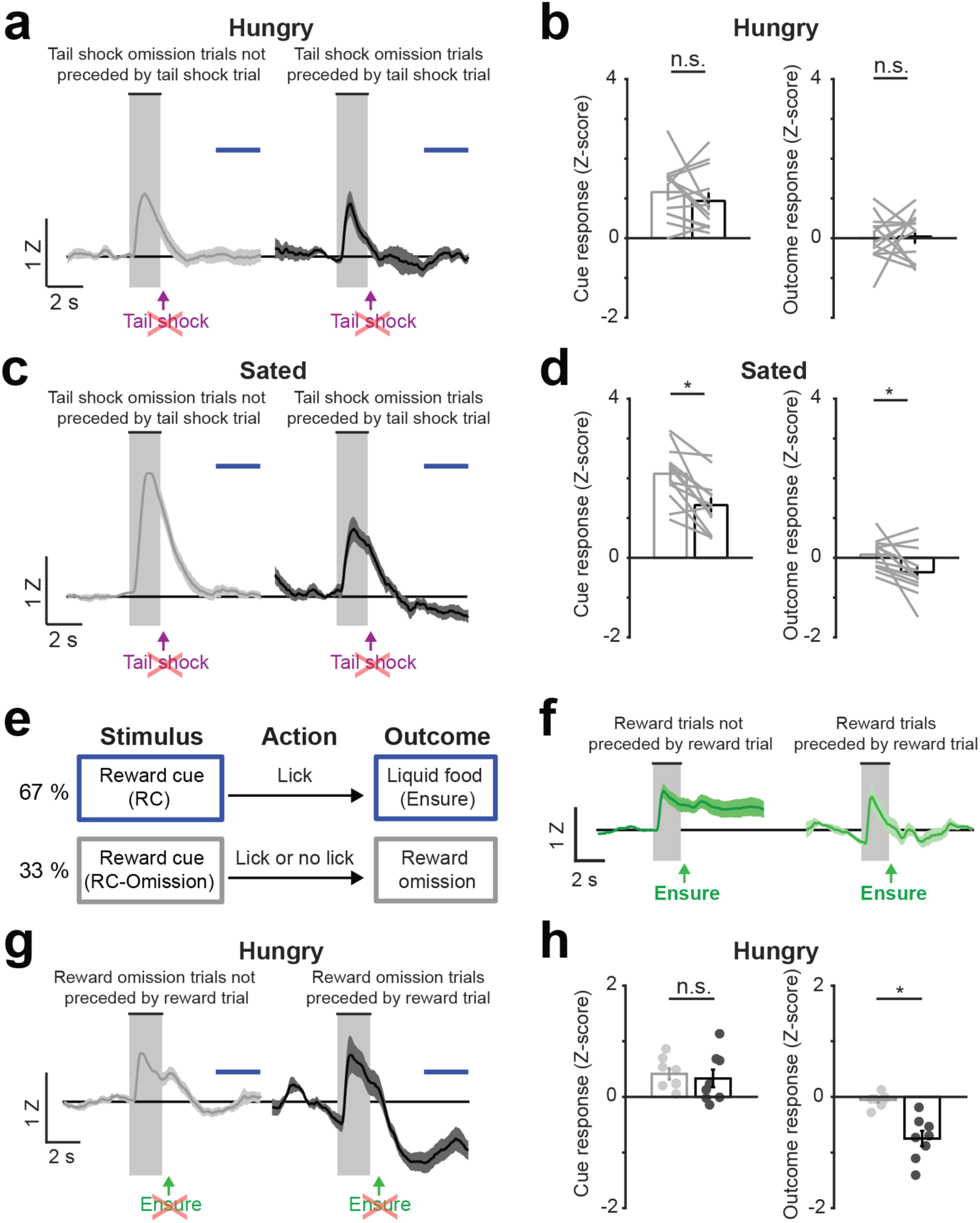
VTA^DA→BA^ axons show an increase in tonic inter-trial activity with reward expectation and a decrease in activity upon reward/shock omission, but only when omission is preceded by a previously rewarded/shocked trial, respectively. **a,** *Left*: mean timecourse for tail shock omission trials that *were not* preceded by trials with tail shock delivery in the hungry state. *Right*: mean timecourse for tail shock omission trials that *were* preceded by trials with tail shock delivery in the hungry state. Error bars: s.e.m. across 13 sessions from 8 mice. The window used for analysis of outcome responses is indicated by a blue bar. **b,** *Left:* cue response magnitude for tail shock omission trials that were not preceded (light gray) or were preceded (dark gray) by tail shock were not significantly different (Wilcoxon sign-rank). *Right:* mean activity after tail shock omission was not different between trials that were or were not preceded by tail shock in hungry mice. Error bars: s.e.m. across 13 sessions from 8 mice. n.s. = not significant. **c,** *Left*: mean timecourse for tail shock omission trials that *were not* preceded by trials with tail shock delivery in the sated state. *Right*: mean timecourse for tail shock omission trials that *were* preceded by trials with tail shock delivery in the sated state. Error bars: s.e.m. across 13 sessions from 8 mice. The window used for analysis of outcome responses is indicated by a blue bar. **d,** *Left:* cue response magnitude for tail shock omission trials that were not preceded by tail shock (light gray) were larger than those that were preceded by tail shock (dark gray) in sated mice. *Right:* mean decrease in activity after tail shock omission was significantly greater when the prior trial contained a tail shock in sated mice, potentially due to increased expectation of additional tail shocks following tail shock receipt in the sated state when mice may be in a more defensive state. Error bars: s.e.m. across 13 sessions from 8 mice. * p < 0.05, Wilcoxon sign-rank. **e,** Modified task design for assessing effects of reward expectation and omission on activity of VTA^DA→BA^ axons. For experiments presented in this Supplementary Figure 8e-h only, we used a modified task that only involved presentation of a single type of cue – the reward cue (RC). Reward cues were followed either by reward delivery (67% of trials) or by reward omission (33% of trials). **f,** *Left:* responses were persistently elevated for many seconds following termination of reward consumption (well beyond the ∼1-s timescale associated with decay of GCaMP6s fluorescence) for trials that followed a *non-rewarded* trial. *Right:* reward trials following a previously *rewarded* trial did not show persistently elevated activity following cue offset (right of the shaded box) relative to pre-cue baseline. Error bars: s.e.m. across 8 mice. Note that response timecourses were re-zeroed to pre-cue baseline. **g,** *Left:* mean timecourse of reward omission trials that *were not* preceded by a rewarded trial. *Right:* mean timecourse of reward omission trials that *were* preceded by a rewarded trial. Error bars: s.e.m. across 8 mice. Note that response timecourses were re-zeroed to pre-cue baseline. The window used for analysis of outcome responses is indicated by a blue bar. **h,** *Left:* mean cue response for reward omission trials that were not preceded (light gray) or were preceded (dark gray) by reward were not significantly different (p > 0.05, Wilcoxon sign-rank). *Right:* mean decrease in activity after reward omission was significantly greater when the prior trial was rewarded, potentially due to increased expectation of additional rewards following reward receipt. Error bars: s.e.m. across 8 mice. * p < 0.05, Wilcoxon sign-rank.

**Supplementary Fig. 9.**
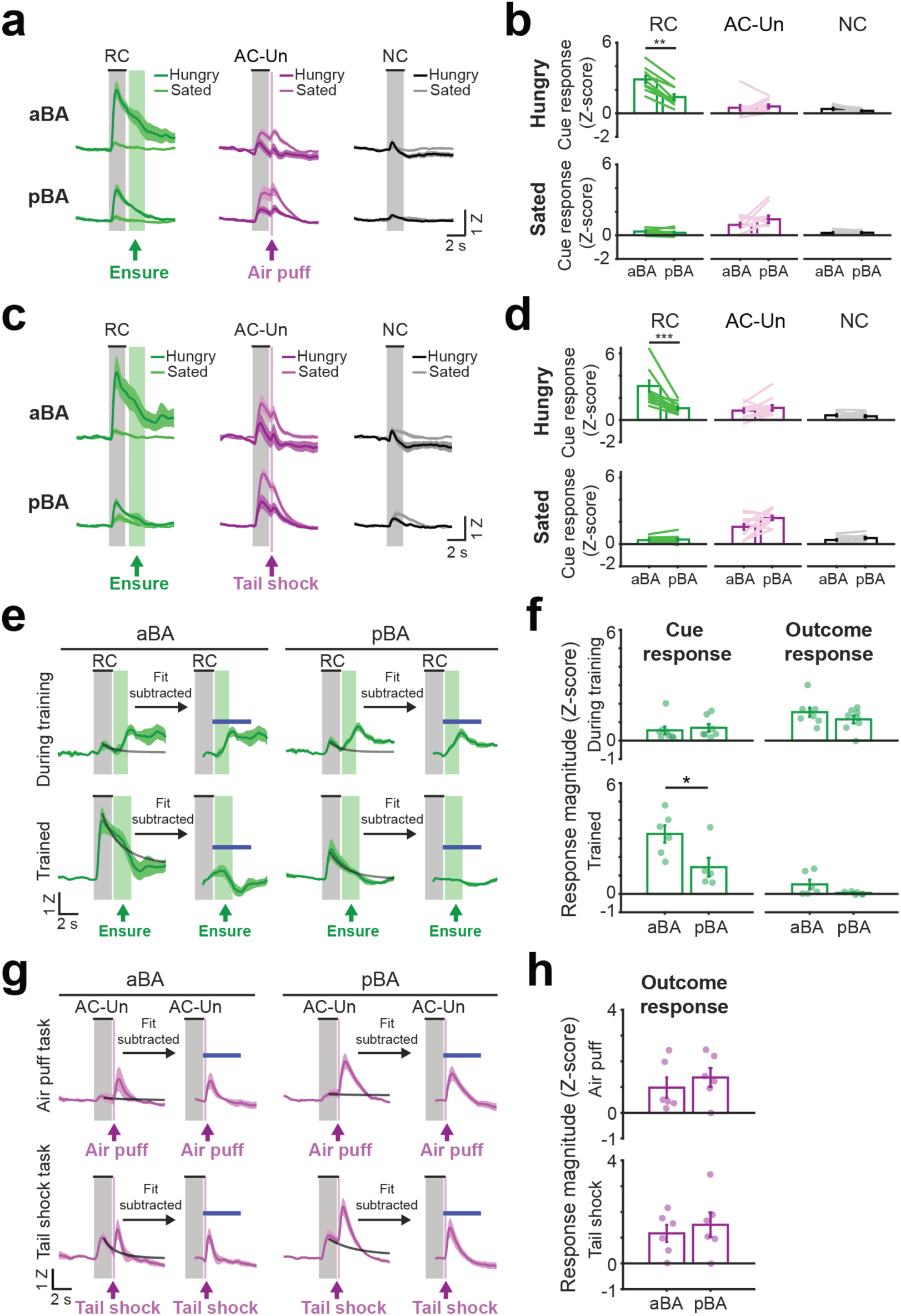
VTA^DA→BA^ axon cue responses are similar across anterior BA and posterior BA. **a,** Mean timecourse during trials involving a reward cue (RC), a neutral cue (NC), or an aversive cue predicting unavoidable air puff (AC-Un), for anterior BA (aBA) and posterior BA (pBA) recordings. Error bars: s.e.m. across 10 sessions from 6 mice. **b,** Cue response magnitude of VTA^DA→BA^ recordings in aBA or pBA (n = 10 sessions from 6 mice). **p < 0.01, Wilcoxon sign-rank. **c,** Mean timecourse during trials involving a reward cue (RC), a neutral cue (NC), or an aversive cue predicting unavoidable tail shock (AC-Un), for aBA and pBA recordings. Error bars: s.e.m. across 11 sessions from 8 mice. **d,** Cue response magnitude of VTA^DA→BA^ axon recordings from anterior vs. posterior BA (n = 11 sessions from 8 mice). *** p < 0.001, Wilcoxon sign-rank. **e,** Mean response timecourses of VTA^DA→BA^ axons during training and following completion of training on avoidable quinine task, from aBA (*left*) or pBA (*right*). Response timecourse of Ensure delivery related activity was obtained by subtracting a monoexponential fit of the cue response. The window used for analysis of Ensure delivery responses is indicated by a blue bar. Error bars: s.e.m. across 8 “during training” and 6 “trained” mice. Z: Z-score. **f,** Cue and outcome response magnitudes of VTA^DA→BA^ axon recordings from aBA or pBA during training (n = 8 mice) vs. following training (n = 6). During training, no differences were found between aBA and pBA (p > 0.05, Wilcoxon rank-sum). Following training, similar to Supplementary Fig 9b,d, RC response magnitudes were greater in aBA. * p < 0.05, Wilcoxon rank-sum. **g,** Mean response timecourses of VTA^DA→BA^ axons during first sessions of training with air puff (top) or tail shock (bottom) in aBA (left) and pBA (right). Response timecourse of aversive outcome delivery-related activity was obtained by subtracting a monoexponential fit of the cue response. The window used for analysis of air puff or tail shock delivery responses is indicated by a blue bar. Error bars: s.e.m. across 6 mice. **h,** Outcome response magnitudes of VTA^DA→BA^ axon recordings from aBA or pBA during first day of air puff training (top) or first day of tail shock training (bottom). Responses to delivery of air puff or tail shock were not different between aBA and pBA.

**Supplementary Fig. 10.**
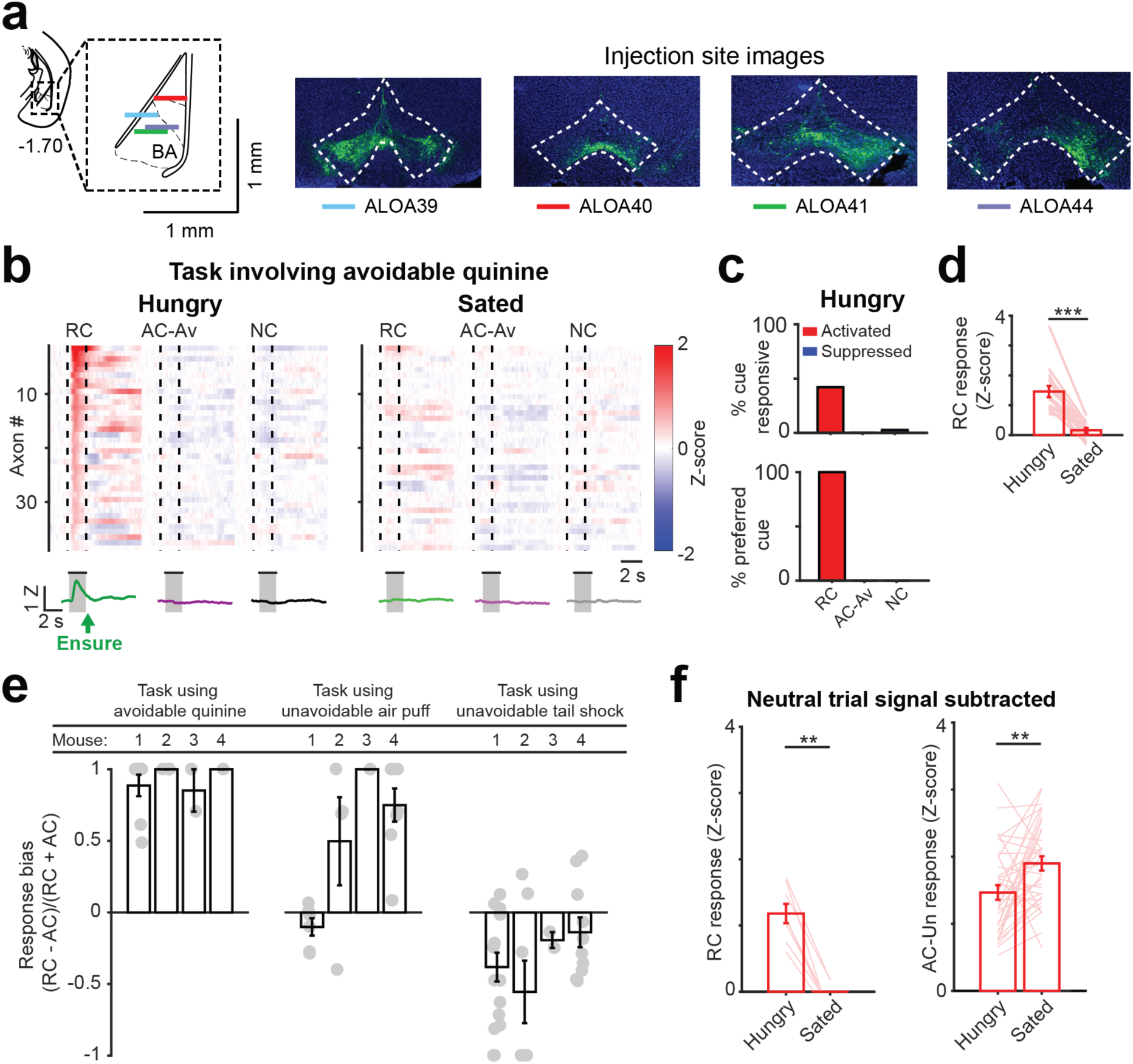
Individual VTA^DA→BA^ axon cue responses are restricted to the reward cue during a task involving aversive cues predicting passively avoidable quinine. **a,** *Left:* location of GRIN lens implants in medial BA (mBA) from VTA^DA→BA^ axon imaging experiments (horizontal colored lines; lenses are depicted at scale), determined from post-hoc histology (n = 4 mice). *Right:* images of injection site showing GCaMP6s-labeled dopamine neurons in VTA from all 4 mice. **b,** *Top*: mean response of individual axons (rows) to presentation of the reward cue (RC), aversive cue predicting avoidable quinine (AC-Av), and neutral cue (NC) (n = 38 axons, 7 fields of view from 4 mice). *Bottom:* population responses (mean ± s.e.m. across 38 axons). **c,** *Top*: percent of all axons with significant cue responses during hungry runs (RC: 16/38 neurons; AC-Av: 0/38; NC: 1/38). *Bottom*: percent of cue responsive axons preferring a given cue. All data are from hungry runs. **d,** Mean RC response for activated axons. Lines: individual axon responses across hunger and satiety (*** p < 0.001, Wilcoxon sign-rank). **e,** Response bias of individual VTA^DA→BA^ axons activated by RC and/or AC from 4 mice following training with avoidable quinine (*left*), following training with air puff (*middle*), and following training with tail shock (*right*). Note that axons recorded from all four mice were initially biased to the RC and shifted towards equal preference of RC and AC following training with air puff and tail shock. **f,** RC (*left*) and AC-Un (*right*) response magnitudes of VTA^DA→BA^ axons during hungry vs. sated states, after subtraction of NC responses. ** p < 0.01, Wilcoxon sign-rank.

**Supplementary Fig. 11.**
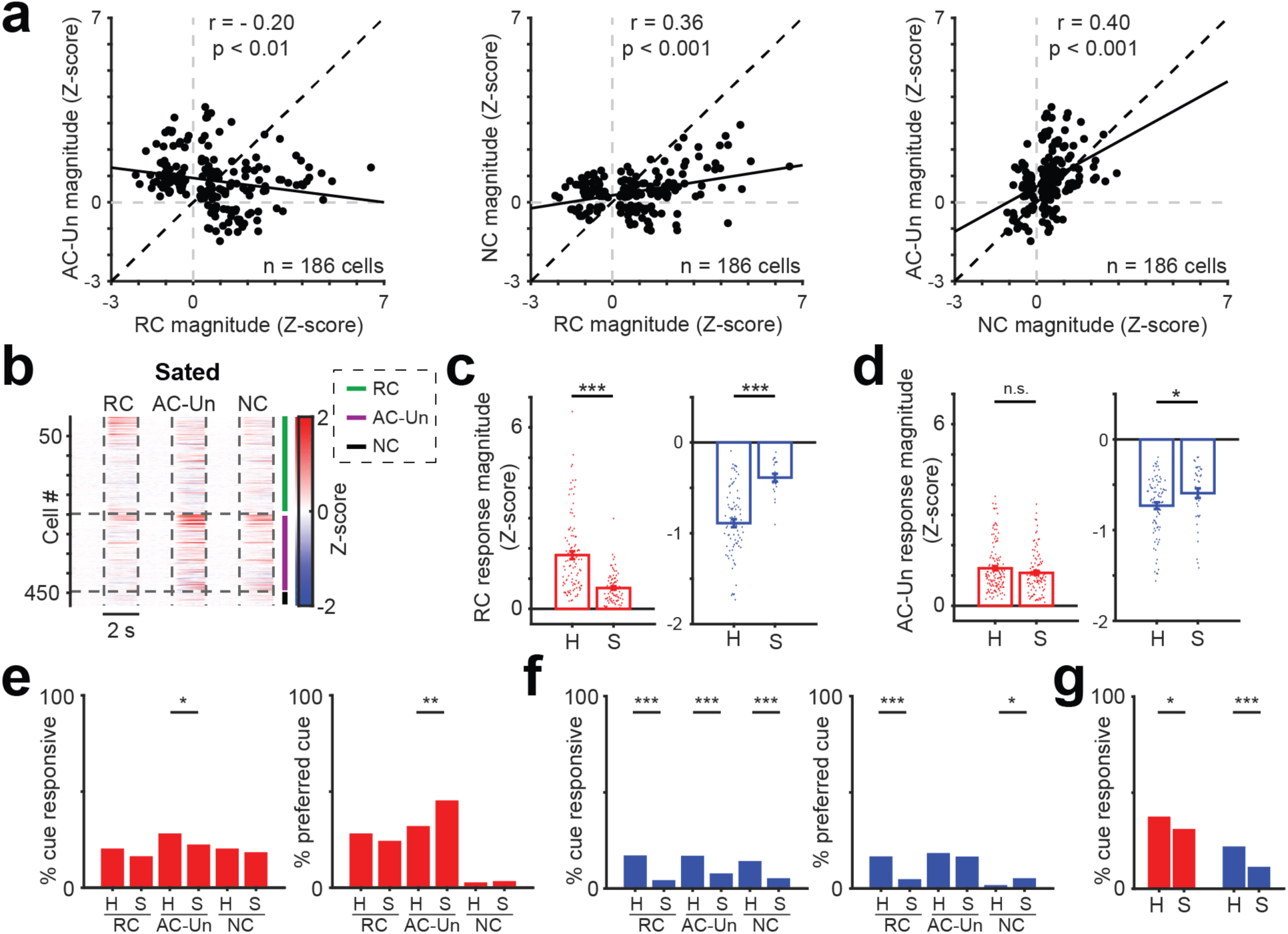
Additional analyses of BA cell body responses during the task involving cues predicting unavoidable tail shock. **a,** *Left:* correlation between magnitude of responses to the reward cue (RC) and to the aversive cue predicting unavoidable tail shock (AC-Un) across basal amygdala (BA) neurons (n = 186 neurons; Pearson’s r: −0.20, p < 0.01). *Middle:* correlation between RC and NC response magnitude (n = 186 neurons; Pearson’s r: 0.36, p < 0.001). *Right:* correlation between NC and AC-Un response magnitude (n = 186 neurons; Pearson’s r: 0.40, p < 0.001). **b,** Heatmap showing mean cue response timecourses of BA neurons (rows) from sated mice. Rows are sorted by response magnitude during earlier runs in the same session, in hungry mice (see Fig. 5c; n = 482 neurons, 9 fields of view from 4 mice). Vertical dashed lines demarcate visual stimulus onsets and offsets. Horizontal lines demarcate sorting of axons by preferred cue (cue with the largest absolute value response). **c,** *Left:* response magnitude of neurons that were significantly activated by the RC during hunger or satiety (n = 98 in hungry mice, n = 79 in sated mice, *** p < 0.001, Wilcoxon rank sum). *Right:* response magnitude of neurons that were significantly suppressed by the RC during hunger or satiety (n = 83 in hungry mice, n = 21 in sated mice, *** p < 0.001, Wilcoxon rank sum). **d,** *Left:* response magnitude of neurons that were significantly activated by the AC-Un during hunger or satiety (n = 136 in hungry mice, n = 108 in sated mice, n.s. = not significant, Wilcoxon rank sum). *Right:* response magnitude of neurons that were significantly suppressed by AC-Un during hunger or satiety (n = 82 in hungry mice, n = 38 in sated mice, n.s. = not significant, p > 0.05, Wilcoxon rank sum). **e,** *Left*: percent of BA neurons that were significantly activated by cues in hungry and sated mice. There was a higher percentage of neurons significantly activated by AC-Un vs. sated mice. *Right*: percent of BA neurons with a given cue preference (activated) in hungry and sated states. There was a higher percentage of BA neurons preferentially activated by the AC-Un in the sated state. * p < 0.05, ** p < 0.01, binomial proportion test. **f,** *Left*: percent of BA neurons that were significantly suppressed by cues in hungry and sated states. There was a higher percentage of suppressed BA neurons in hungry mice. *Right*: percent of BA neurons with a given cue preference (suppressed) in hungry and sated states. There was a higher percentage of BA neurons preferentially suppressed by the RC in the hungry state and NC in the sated state. * p < 0.05, *** p < 0.001, binomial proportion test. **g,** Percent of BA neurons significantly activated (*left*) or suppressed (*right*) by any cue in hungry vs sated states. * p < 0.05, *** p < 0.001, binomial proportion test.

**Supplementary Fig. 12.**
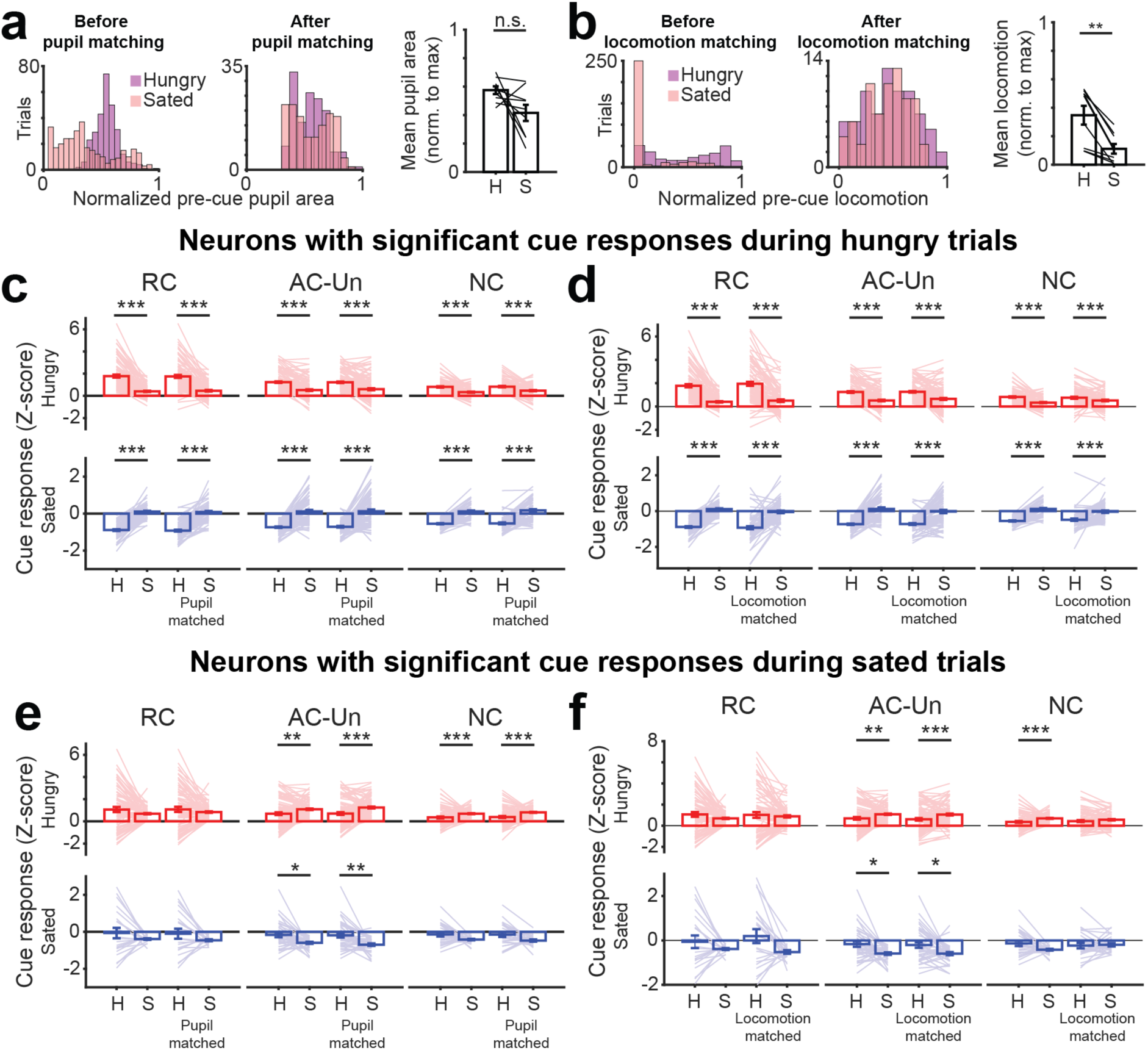
Changes in pupil area and locomotion across states do not account for observed changes in BA activity in mice trained with cue predicting rewards and cues predicting unavoidable tail shock. **a,** Histograms of pupil areas in the 2 s preceding cues during hungry and sated states from an example session, before (*left*) and after (*right*) matching trials for pupil area (see Methods). *Right:* mean pupil area in the 2 s preceding a cue during hungry and sated states (normalized to maximum pupil area during that entire session across both states). Mean ± s.e.m. across 9 sessions from 4 mice. n.s.= not significant, Wilcoxon sign-rank. **b,** Histograms of locomotion in the 2 s preceding cues during hungry and sated states from an example session before (*left*) and after (*right*) matching trials for locomotion. *Right:* mean locomotion in the 2 s preceding a cue during hungry and sated states (normalized to maximum locomotion during that entire session across both states). Mean ± s.e.m. across 9 sessions from 4 mice. ** p < 0.01, Wilcoxon sign-rank. **c,** Comparison of cue responses of BA neurons that were significantly responsive in the hungry state. Attenuation of cue responses following satiation persisted after matching pupil area distributions across states. *** p < 0.001, Wilcoxon sign-rank. **d,** Comparison of cue responses of BA neurons that were significantly responsive in the hungry state. Attenuation of cue responses following satiation persisted after matching locomotion distributions across states. *** p < 0.001, Wilcoxon sign-rank. **e,** Comparison of cue responses of BA neurons that were significantly responsive in the sated state. Enhancement of cue responses following satiation persisted after matching pupil area distributions across states. * p < 0.05, ** p < 0.01, *** p < 0.001, Wilcoxon sign-rank. **f,** Comparison of cue responses of BA neurons that were significantly responsive in the sated state. Enhancements of cue responses following satiation persisted after matching locomotion distributions across states. * p < 0.05, ** p < 0.01, *** p < 0.001, Wilcoxon sign-rank.

**Supplementary Fig. 13.**
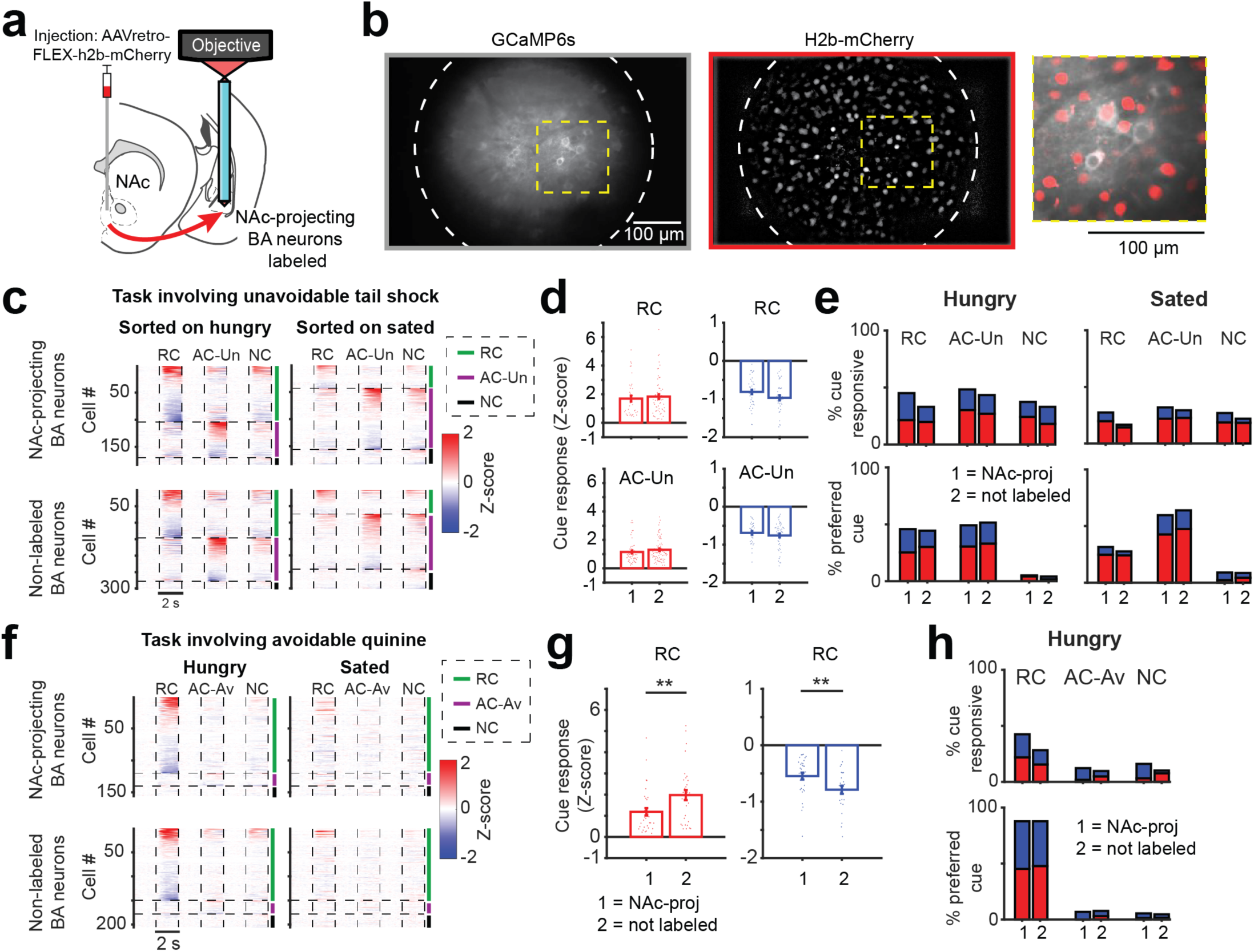
Similar findings in NAc-projecting BA neurons as in nearby unlabeled BA neurons. **a,** Schematic depicting retrograde labeling of nucleus accumbens (NAc)-projecting basal amygdala (BA) neurons and two-photon imaging of these labeled neurons (see Methods for additional details). **b,** *Left top:* mean image of GCaMP6s fluorescence expression in excitatory BA neurons in an example field of view. *Left bottom:* mean image from the same field of view as above, showing mCherry fluorescence in the nuclei of retrogradely labeled NAc-projecting BA neurons. *Right:* overlaying the GCaMP6s and mCherry fluorescence within the yellow box allows identification of neurons with and without red-labeled nuclei. **c,** *Top left:* heatmap with rows depicting mean responses of NAc-projecting BA neurons (n = 182 neurons, 9 fields of view from 4 mice) from the task involving an aversive cue predicting unavoidable tail shock (AC-Un). *Top right:* responses of the same neurons as in *top left,* but during sated trials. In this panel, neurons (rows) are sorted by response magnitude during the sated state, to facilitate comparison of the two subpopulations of BA neurons. *Bottom left:* heatmap with rows depicting mean responses of BA neurons that are not retrogradely labeled with nuclear-localized mCherry (n = 300 neurons, 9 fields of view from 4 mice), from the task involving unavoidable tail shock. *Bottom right:* responses of the same neurons as in *bottom left* but during sated trials and sorted by response magnitude in the sated state. Note the similarity in response profiles across the two subpopulations. These data, combined across NAc-projecting BA neurons and other BA neurons, are also shown in Figure 6c. **d,** *Top left:* Response magnitude was not significantly different in NAc-projecting BA neurons activated by the RC (‘1’, n = 39) vs. non-labeled BA neurons activated by the RC (‘2’, n = 59) (p > 0.05, Wilcoxon rank sum). *Top right:* similarly, response magnitude was not significantly different in NAc-projecting BA neurons suppressed by the RC (‘1’, n = 43) vs. non-labeled BA neurons suppressed by the RC (‘2’, n = 40) (p > 0.05, Wilcoxon rank sum). *Bottom left:* response magnitude was not significantly different in NAc-projecting BA neurons activated by the AC-Un (‘1’, n = 55) vs. non-labeled BA neurons activated by the AC-Un (‘2’, n = 81) (p > 0.05, Wilcoxon rank sum). *Bottom right:* similarly, response magnitude was not significantly different in NAc-projecting BA neurons suppressed by the AC-Un (‘1’, n = 33) vs. non-labeled BA neurons suppressed by the AC-Un (‘2’, n = 49) (p > 0.05, Wilcoxon rank sum). **e,** *Top left:* percent of significantly responsive neurons from either NAc-projecting BA neurons (‘1’) and for non-labeled BA neurons (‘2’) during hungry trials. *Top right:* percent of significantly responsive neurons from either NAc-projecting BA neurons (‘1’) or from non-labeled neurons (‘2’) during sated trials. *Bottom left:* percent of neurons with a given cue preference of NAc-projecting BA neurons (‘1’) or from non-labeled neurons (‘2’) during hungry trials. *Bottom right:* percent of neurons with a given cue preference for NAc-projecting BA neurons (‘1’) vs. non-labeled BA neurons (‘2’) during hungry trials. Red: activated neurons. Blue: suppressed neurons. **f,** *Top left:* heatmap with rows depicting mean responses of NAc-projecting BA neurons (n = 155 neurons, 15 fields of view from 7 mice) from task using avoidable quinine (AC-Av). *Top right:* responses of the same neurons with same sorting as in *top left,* but during sated trials. *Bottom left:* rows depicting mean responses of BA neurons that lack nuclear mCherry labeling (n = 205 neurons, 15 fields of view from 7 mice), from the task involving avoidable quinine. *Bottom right:* responses of the same neurons with the same sorting as in *bottom left* but during sated trials. Note: unsorted data combined from NAc-projecting and other BA neurons also shown in Figure 1h. g, *Left:* for the above task involving reward cues (RC) and cues predicting passively avoidable quinine (AC-Av), RC response magnitude for NAc-projecting BA neurons (‘1’, n = 34) that were activated by cue presentation (red) showed significantly weaker response magnitudes compared to non-labeled BA neurons (‘2’, n = 32). ** p < 0.01, Wilcoxon rank sum. *Right:* similarly, RC response magnitude for NAc-projecting BA neurons (‘1’, n = 32) suppressed by cue presentation (blue) showed significantly weaker response magnitudes compared to non-labeled BA neurons (‘2’, n = 26). ** p < 0.01, Wilcoxon rank sum. Mean ± s.e.m. across neurons. **h,** *Top:* percent of significantly responsive NAc-projecting BA neurons (‘1’) or significantly responsive non-labeled BA neurons (‘2’). *Bottom:* cue preference of significantly responsive NAc-projecting BA neurons (‘1’) and of significantly responsive, non-labeled BA neurons (‘2’). Red: activated neurons. Blue: suppressed neurons.

**Supplementary Fig. 14.**
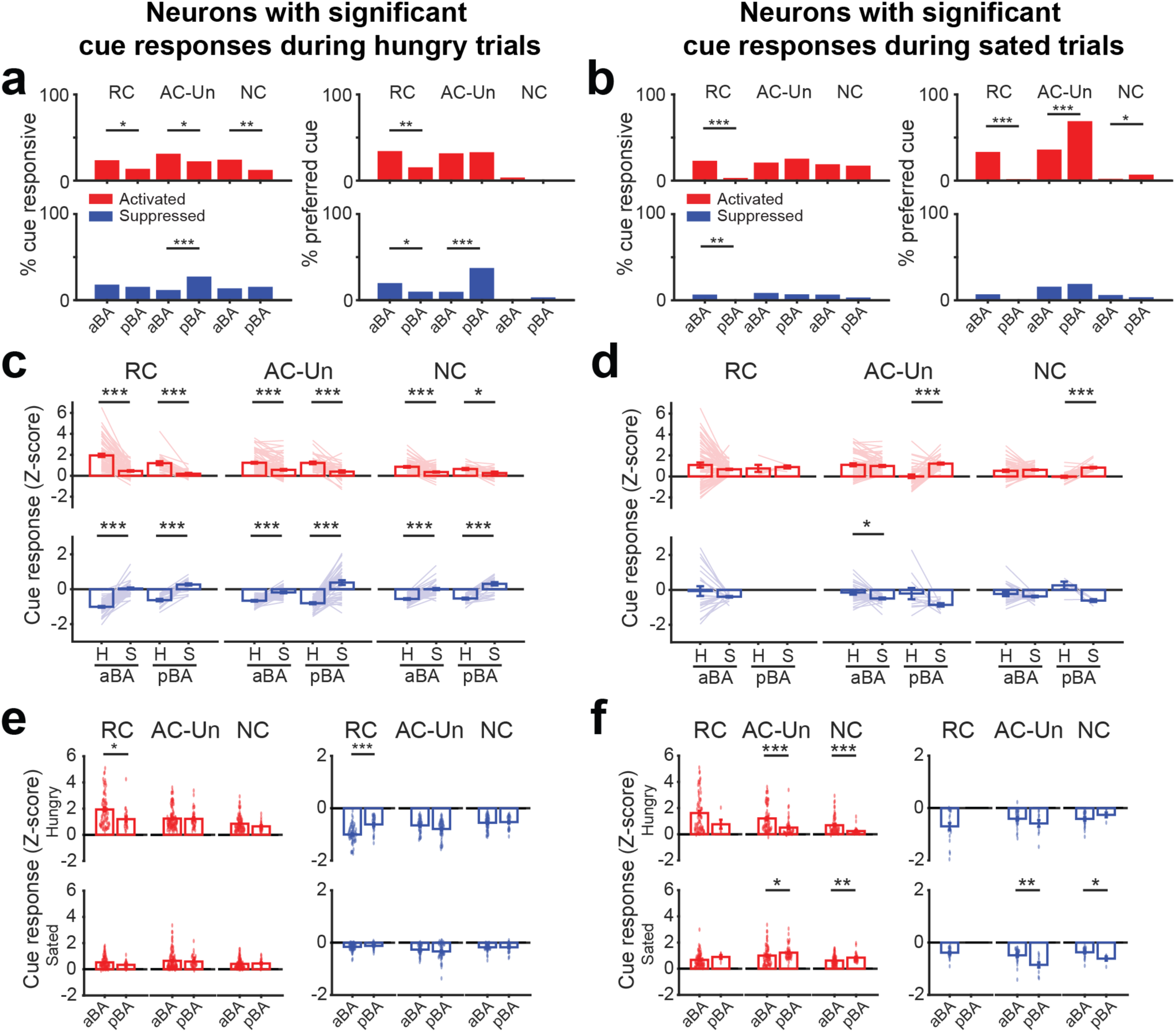
Analyses of BA cell body cue responses in anterior and posterior subregions of BA. **a,** Percent of BA neurons in anterior BA (aBA) vs. posterior BA (pBA) that were significantly activated (*top left*) or suppressed (*bottom left*) in hungry mice by RC, AC-Un, or NC. There was a higher percentage of aBA neurons activated by each of the three cues compared to pBA neurons, whereas there was a higher percentage of pBA neurons suppressed by the AC-Un. Percent of BA neurons in aBA vs. pBA that were preferentially activated (*top right*) or suppressed (*bottom right*) by cues in hungry mice. The percentage of aBA neurons that were preferentially activated by the RC was higher than in pBA whereas the percentage of pBA neurons that were preferentially suppressed by the AC-Un was higher in pBA. * p < 0.05, ** p < 0.01, *** p < 0.001, binomial proportion test. **b,** *Left*: percent of BA neurons in anterior BA (aBA) vs posterior BA (pBA) that were significantly activated (*top left*) or suppressed (*bottom left*) in sated mice by RC, AC-Un, or NC. There was a higher percentage of aBA neurons activated by the RC and suppressed by the RC. *Right:* percent of BA neurons in aBA vs. pBA that were preferentially activated (*top right*) or suppressed (*bottom right*) by cues in sated mice. The percentage of aBA neurons that were preferentially activated by the RC was higher than for pBA neurons, whereas the percentage of pBA neurons that were preferentially activated by the AC-Un was higher for pBA neurons. * p < 0.05, ** p < 0.01, *** p < 0.001, binomial proportion test. **c,** Comparison of cue responses of aBA and pBA neurons that were significantly responsive in the hungry vs. sated states. Attenuation of cue response magnitudes following satiation was observed in both aBA and pBA. * p < 0.05, *** p < 0.001, Wilcoxon sign-rank. **d,** Comparison of cue responses of aBA and pBA neurons that were significantly responsive in hungry vs. sated states. Enhancement of the AC-Un and NC response magnitudes following satiation was observed in pBA, but not in aBA. * p < 0.05, *** p < 0.001, Wilcoxon sign-rank. **e,** Comparison of cue response magnitudes of aBA vs. pBA (*left:* activated; *right:* suppressed) neurons which were found to be responsive in the hungry state. The RC response was greater in aBA in hungry mice. * p < 0.05, *** p < 0.001, Wilcoxon sign-rank. **f,** Comparison of cue response magnitudes of aBA vs. pBA (*left:* activated; *right:* suppressed) neurons which were found to be responsive in the sated state. The AC-Un and NC responses were greater in aBA in hungry mice whereas the AC-Un response was greater in pBA in sated mice. * p < 0.05, ** p < 0.01, *** p < 0.001, Wilcoxon sign-rank.

**Supplementary Fig. 15.**
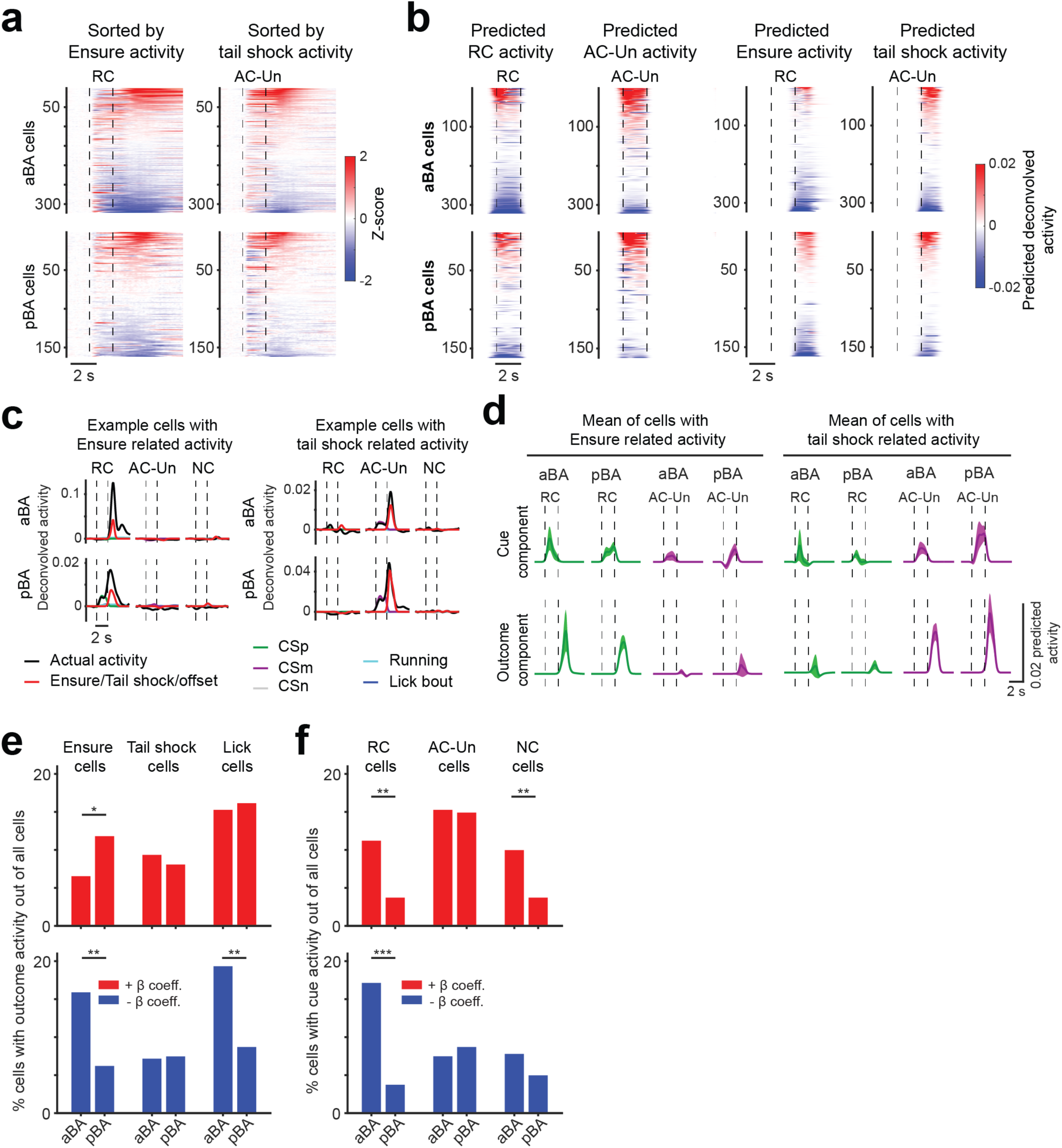
Analyses of cell body outcome responses in anterior and posterior subregions of BA using a generalized linear model to estimate distinct components of overall activity. **a,** Heatmap showing mean cue response timecourses of BA neurons (rows) from anterior BA (aBA; *top*) and from posterior BA (pBA; *bottom*) during hungry sessions, sorted by response magnitude in the 2 s following cue offset (n = 321 aBA neurons, 5 fields of view from 2 mice; n = 161 pBA neurons, 4 fields of view from 2 mice), for the task involving unavoidable tail shock. Vertical dashed lines demarcate visual stimulus onsets and offsets. **b,** *Left:* heatmap showing mean generalized linear model (GLM) predicted cue-related activity for RC and AC-Un from aBA (*top*) and pBA (*bottom*) neurons, sorted by response magnitude during 2 s cue period (see Methods). *Right:* heatmap showing mean GLM Ensure/tail shock-related activity from aBA (*top*) and pBA (*bottom*) neurons, sorted by response magnitude during 2 s following cue offset. **c,** *Left:* example neurons from aBA (*top*) and pBA (*bottom*) which had significant (see Methods) Ensure-related activity. *Right:* example neurons from aBA (*top*) and pBA (*bottom*) which had significant tail shock-related activity. Actual deconvolved neuronal activity (black lines), Ensure-related activity (red lines) (RC trials)/tail shock(AC-Un trials)/offset(NC trials), cue-related activity (green: RC; purple: AC-Un; gray: NC), running-related activity (cyan lines), and lick bout-related activity (blue lines). **d,** Mean predicted activity timecourses for cues (*top*) and outcomes (*bottom*) using all GLM components, from all cells with significant Ensure (*left*) or tail shock (*right*) components. Note that cells with Ensure components preferentially exhibited RC-related responses and cells with tail shock components preferentially exhibited AC-Un-related responses. **e,** Percent of all aBA vs. pBA neurons that have significant predicted activity related to Ensure (*left*), tail shock (*middle*), or licking (*right*). *Top:* neurons with positive beta-coefficients indicative of predicted increases in activity. *Bottom:* neurons with negative beta-coefficients. There was a larger percentage of neurons in pBA with significant predicted increases in activity to Ensure while there was a larger percentage of cells with Ensure- and Lick-related decreases in activity in aBA vs. pBA. * p < 0.05, ** p < 0.01, binomial proportion test. **f,** Percent of all aBA vs. pBA neurons that have significant activity related to presentation of the RC (*left*), AC-Un (*middle*), or NC (*right*). Similar to results in Supplementary Fig. 14, there was a larger percentage of neurons in aBA with significant predicted increases and predicted suppression in activity related to the RC, and with with significant predicted increases in activity related to the NC. ** p < 0.01, *** p < 0.001, binomial proportion test.

**Supplementary Movie 1 | Movies of average VTA^DA→BA^ axon responses to salient and non-salient cues**

*Left:* movie of cue-evoked responses, averaged across all reward cue (RC) trials (n = 58 trials) from a single session. *Middle:* movie of cue-evoked responses, averaged across all presentations of the aversive cue predicting unavoidable air puff (AC-Un) (n = 53 trials). *Right:* movie of cue-evoked responses, averaged across all presentations of the neutral cue (NC) (n = 56 trials). Movies have been spatially down-sampled by 2x and temporally binned by 4 frames. Frame rate: 7 frames/s (i.e. movie is playing at 1.8x real-time). Stimulus duration (indicated by white squares) was two seconds. White circles indicate perimeter of field of view. Percent change in GCaMP6s fluorescence ranges from −15% (black) to +15**%** (white). Gray: no change in activity. See also Fig. 5b.

